# A multi-omics and cell type-specific characterization of the ventral striatum in human cocaine use disorder

**DOI:** 10.1101/2024.10.09.617337

**Authors:** Eric Zillich, Annasara Artioli, Andrea C. Rossetti, Diana Avetyan, Hanna Belschner, Josef Frank, Frank Stein, Jennifer J. Schwarz, Naguib Mechawar, Gustavo Turecki, Markus M. Nöthen, Anita C. Hansson, Christian C. Witt, Marcella Rietschel, Philipp Koch, Rainer Spanagel, Lea Zillich, Stephanie H. Witt

**Author notes:** co-senior authorship (Lea Zillich, Stephanie H. Witt).

## Abstract

Epigenome, transcriptome, and proteome analyses of postmortem brains have revealed initial molecular insights into cocaine use disorder (CUD). However, the inter-relationship between these –omics and the contribution of individual cell types remain largely unknown. We present an in-depth analysis of molecular changes in the ventral striatum in CUD at multi-omics and single-cell resolution. Integrative multi-omics analyses of microRNA-seq, RNA-seq, and proteomics datasets in 41 individuals and single-nuclei RNA-seq in a subset of 16 individuals revealed conserved deregulation of metabolic pathways, oxidative phosphorylation, and glutamatergic signaling. Cell type-specific analyses identified inverse metabolic pathway deregulation patterns in glial and neuronal cells, notably in astrocytes and medium spiny neurons (MSNs). Characterizing astrocyte-neuron crosstalk revealed altered glutamatergic and cell adhesion signaling in CUD. By applying a comprehensive multi-omics analytical framework, our study provides novel insights into CUD-associated molecular changes in the ventral striatum, suggesting astrocytes, MSNs, and their crosstalk as particularly perturbed in CUD.

## Introduction

Individuals with cocaine use disorder (CUD) present with an excessive intake of the psychostimulant cocaine despite negative consequences, strong cocaine craving, and relapse after periods of abstinence^1^. In 2021, a total of 21.6 million individuals have used cocaine worldwide^2^ with around 20% of cocaine users transitioning from episodic cocaine use to use disorder during their lifetime^3^. Current treatment options are limited and retrospective analyses suggest that less than 25% of treatment-seeking patients remain abstinent after completing an inpatient treatment program^4^. A deeper understanding of the neurobiological mechanisms of CUD is essential to provide the basis for the development of mechanism-based interventions.

The brain is assumed to be the most prominently affected organ in the development and maintenance of CUD. Neuroimaging studies revealed structural brain alterations such as reduced gray matter volume in CUD and functional changes in neurocircuit connectivity between different brain regions of the reward system have been described^5–11^. Cocaine has strong neurovascular effects in the brain, it reduces cerebral blood flow and by its direct effect on brain perfusion contributes to neurocircuit alterations^12^. Further, cocaine-induced epigenetic and transcriptional changes in the human brain were proposed as a molecular mechanism involved in the formation of structural and functional neurocircuit changes in individuals with CUD^13^. Investigating molecular signatures of CUD in the human brain thus depicts an important approach toward a better understanding of the underlying disease processes^14^. Previous studies in postmortem human brain tissue have reported on molecular alterations of the epigenome, transcriptome, and proteome in CUD^15–22^. Epigenome-wide studies have so far mainly focused on DNA methylation and showed CUD-associated differential methylation in multiple addiction-relevant brain regions such as the prefrontal cortex (PFC)^15^, the ventral striatum (VS)^17^, and the caudate nucleus (CN)^16^. Differential methylation levels were detected in genes involved in dopamine metabolism such as tyrosine hydroxylase^17^ while at the pathway level, epigenetic changes were related to transcription factor activity and synaptic signaling^15,17^. Another domain of epigenetic regulators are micro-RNAs (miRNAs), small RNA molecules that bind to complementary nucleotide sequences on mRNAs thereby regulating mRNA degradation and translation rate to proteins^23^. While associations with CUD were identified in peripheral blood for miRNAs such as miR-124 and miR-184^18^, differential miRNA expression remains understudied in the human CUD brain. At the transcriptomic scale, multiple studies have identified CUD-associated changes in RNA levels in cortical^21^, limbic^19^, and striatal brain regions^20^. The most prominent findings include alterations of transcripts and co-expression networks involved in neuroplasticity, neuroinflammation, and mitochondrial respiration^19–21^. Further, investigating the proteome is particularly important for evaluating altered neurobiological functions in CUD as protein-protein interactions depict a key component of cellular signaling. Proteomic analysis of the human prefrontal cortex in CUD revealed differential expression of proteins involved in neuroinflammation and myelination supporting the neuroimaging findings of white matter deficits in postmortem brain^22^.

While single-omics studies are valuable in characterizing disease-associated changes in a class of biological molecules such as RNAs or proteins, it remains unclear to what extent these molecular alterations are conserved across layers of biological regulation. Integrative multi-omics analysis of epigenomic regulation, the transcriptome, and the proteome addresses the inter-regulated nature of biological processes thereby depicting a powerful tool to uncover molecular mechanisms of biological deregulation. Another major limitation of individual omics-wide association studies is that cell type-specific associations cannot be sufficiently deduced from bulk-level results increasing the need for analyses at single-cell resolution. First single-nuclei RNA-sequencing (snRNA-seq) studies have been performed in different brain regions mainly focusing on rodent models of CUD^24–27^. While previous findings from bulk transcriptomic studies such as differential expression of neuroplasticity genes were confirmed in snRNA-seq approaches, results were strongly cell type dependent highlighting the importance of further analyses at single cell resolution.

In the present study, we addressed the limitations of single-omics association studies by performing an integrative multi-omics analysis of miRNA-seq, RNA-seq, and proteomic data from the same postmortem brain tissue cohort followed by a cell type-specific investigation of transcriptomic signatures. Analyses were performed in a collection of N=41 postmortem human brain samples of the VS, an important brain region of the neurocircuitry of addiction involved in reward and reinforcement processing^28^. We performed bulk-level multi-omics analyses with two main objectives: the identification of CUD associations at multiple individual molecular levels and the investigation of the inter-relationship of findings across layers of biological regulation. We additionally performed snRNA-seq to evaluate the cell type specificity of transcriptomic changes in CUD. We finally integrated our human snRNA-seq CUD dataset with rodent snRNA-seq data to identify potential converging evidence between human CUD and a controlled cocaine exposure study in rats. Here, we present an in-depth characterization of CUD in postmortem human brain at both bulk and single-nuclei resolution identifying metabolic, synaptic, and immunological changes as molecular hallmarks of the ventral striatum in CUD. Our study further suggests striatal astrocytes and medium spiny neurons as particularly perturbed cell types in CUD.

## Results

From a collection of N=41 postmortem human brain tissue samples (N=20 individuals with CUD, N=21 without CUD) we generated miRNA-seq (N=40), RNA-seq (N=40), proteomics (N=40), and snRNA-seq data (N=16). A graphical summary of the study design and analyses is provided in Figure 1A+B. Dataset availability for each of the VS samples is shown in Figure S1A.

**Figure 1.**
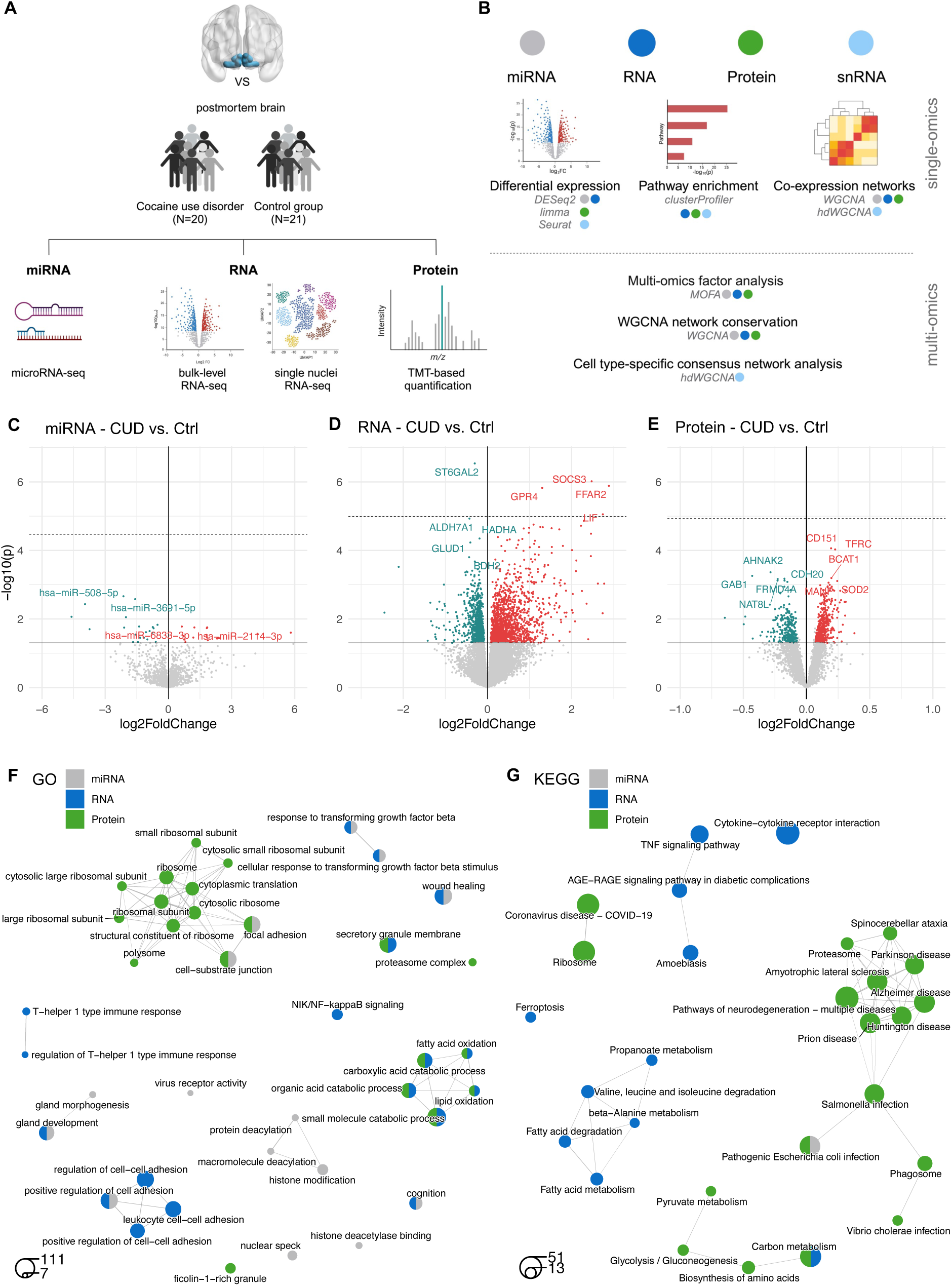
Multi-omics characterization of miRNA, RNA, and protein changes in the ventral striatum in cocaine use disorder. **A** graphical summary of the the study design based on N=20 with and N=21 individuals without cocaine use disorder (CUD/Ctrl). Bulk-level profiling of miRNA (miRNA-seq), RNA (RNA-seq) and protein (TMT-labeling proteomics) changes in CUD and additional snRNA-seq to evaluate the cell type specificity of transcriptomic changes. **B** summary of analysis methods used throughout the present study including individual-level analyses (single-omics) and integrative multi-omics approaches (multi-omics). Bulk-level analysis results of **C** miRNA, **D** RNA, and **E** protein differential expression (DE) analyses. Dashed line indicates 5%-FDR significance, |log2FC| >0.07 (5% change in expression levels). **F** semantic clustering of statistically significant (FDR q<0.05) results from GO enrichment analysis suggests unique and shared biological pathways for differentially expressed genes (p_nominal_<0.05) at the miRNA target gene (grey), RNA (blue) and protein levels (green). **G** results of the same analysis approach for KEGG pathways.

To assess potential systematic differences in phenotypes affecting miRNA, RNA, or protein levels, we evaluated donor demographics. Besides the cause of death, with CUD cases having a significantly higher rate of suicide (p=0.002, Table S1), no significant phenotype differences were found between CUD cases and controls. After PCA-based quality control, 38 individuals remained for miRNA-seq, RNA-seq, and proteomic data analyses (Figure S1B). To address potential confounding from differential cell type fractions, we performed cell type deconvolution using CIBERSORT in the bulk RNA-seq dataset (Figure S1C+D, STAR Methods). This confirmed no significant differences in cell type proportions between the 17 CUD cases and 21 controls (Table S2).

### Individual analysis of bulk-level miRNA-seq, RNA-seq, and proteomic data in the ventral striatum suggests metabolic and immunological alterations in cocaine use disorder

We first performed individual-level differential expression analyses of miRNA-seq, RNA-seq, and proteomics datasets (N=38 individuals) to identify associations with CUD for each of the three molecular levels. Analysis of miRNA-seq data revealed 16 upregulated and 21 downregulated miRNAs in CUD at nominal significance (p<0.05). The strongest association for upregulated miRNAs was found for hsa-miR-6833-3p (log2FC=0.64, p=1.67e-02, q=1), while hsa-miR-508-5p (log2FC=-2.13, p=2.18e-03, q=1) was the top finding among downregulated miRNAs in CUD (Figure 1C, Table S3a). The transcriptome (RNA-seq) analysis identified 36 transcriptome-wide significant differentially expressed genes (DEGs, 5% FDR, Figure 1D, Table S3b). Here, cytokine signaling regulator *SOCS3* (log2FC=2.47, p=9.63e-07, q=7.38e-03) and sialyltransferase *ST6GAL2* (log2FC=-0.30, p=2.90e-07, q=5.71e-03) were the top findings of up- and downregulated DEGs, respectively. Using Tandem Mass Tag (TMT)-based proteomics, we detected 765 differentially expressed proteins (DEPs) in CUD with surface glycoprotein CD151 (log2FC=0.19, p=8.68e-05, q=0.20) and AHNAK2 involved in calcium signaling (log2FC=-0.28, p=4.33e-04, q=0.20) as top up- and downregulated DEPs, respectively (Figure 1E, Table S3c). Of the 765 DEPs, a total of 469 were up- and 296 were downregulated at nominal significance (p<0.05), while no DEP remained statistically significant after multiple testing correction.

To characterize altered biological pathways in the VS, we analyzed CUD-associated nominally significant DEGs and DEPs (p<0.05) in pathway enrichment analyses using Gene Ontology (GO) and KEGG databases as reference (Table S4). Information from the miRNA dataset was added by predicting RNA target genes for the N=37 nominally significant miRNAs (p<0.05) using miRNet^29^ (Table S3a). Here, we detected N=660 predicted RNA targets among the N=2,534 nominally significant DEGs (Table S3b). When we performed clustering of significant GO terms based on biological similarity, we detected a functional GO term module related to fatty acid metabolism that was conserved among RNA and protein results (Figure 1F) supporting previous findings of fatty acid metabolism changes in the brain in a rodent model of cocaine addiction^30^. Additional functional modules were detected that were either RNA- or protein-specific. Here, RNA-specific changes were detected in cell adhesion and immunological signaling while protein-specific enrichment was related to ribosomal pathways. These pathway modules were also significantly enriched for miRNA target DEGs suggesting miRNA-mediated deregulation as a contributing factor to cell adhesion and immunological alterations in CUD. KEGG pathway analysis confirmed the overrepresentation of DEGs and DEPs within metabolic, ribosomal, and immunological processes (Figure 1G). Further, an additional DEP-specific module was found to be associated with neurodegenerative diseases that show overlapping symptoms with CUD, such as brain atrophy^31^.

To further elaborate on the relationship between transcriptomic and proteomic profiles of the VS, we performed a transcriptome-proteome correlation analysis using expression information from 3,935 genes for which both, RNA and protein data were available (see STAR Methods). Based on mean RNA and protein expression levels across samples, we observed a moderate Pearson correlation of *r*=0.43 (p<2.2e-16) between transcriptome and proteome (Figure S2A-C, Table S5) well in line with results from another transcriptome-proteome correlation analysis by Luo et al. in prefrontal cortex^32^. In addition, we found the correlation to be independent of CUD status (Figure S2D). To identify genes for which RNA levels predict protein levels particularly well or poorly, we selected the genes with the strongest positive and negative correlation of RNA and protein levels in the VS. Here, strong concordance of RNA and protein levels was observed for synaptic signaling genes while an inverse relationship between RNA and protein levels was most prominent among oxidative phosphorylation genes (Figure S2E+F).

### Proteo-transcriptomic co-expression network analysis reveals conservation of CUD-associated metabolic gene networks across RNA and protein levels

Profiling gene networks allows the identification of co-regulated gene expression programs that often provide more information about altered biological processes than deregulation patterns of individual genes. To construct co-expression modules at the multi-omics scale we first performed weighted correlation network analyses (WGCNA) in miRNA, mRNA, and protein datasets individually and then performed integrative analysis of networks across -omics. In the miRNA expression dataset, we found 8 co-expression modules (Figure S3A) for which no significant association with CUD was detected (Figure S3B). Network construction in RNA-seq data revealed 54 co-expression modules of which 4 (“grey60”, “lightcyan1”, “skyblue3” and “midnightblue”) were significantly associated with CUD (p<0.05, Figure 2A, Figure S3C, Table S6). RNA module “grey60” (*r*=-0.45, p=0.004) was enriched for astrocytic and fatty acid metabolism pathways and a significant positive association with astrocytes was found (*r*=0.46, p=0.004). Module “lightcyan1” (*r*=0.34, p=0.04) consisted of genes overrepresented in chromatin remodeling pathways, while “skyblue3” was associated with aldehyde dehydrogenase and oxidoreductase activity (Table S7, Figure 2C). In the proteomic dataset, we found 23 protein co-expression modules of which 3 displayed a significant association with CUD including modules “yellow”, “brown” and “tan” (Figure 2B, Figure S3D, Table S6). Protein modules “yellow” and “brown” displayed significant enrichment for pathways previously identified at the RNA level including fatty acid metabolism (“yellow”+“brown”) and oxidoreductase activity (“brown”, Figure 2C, Table S7). Biological functions unique to the protein level were pathways involved in synaptic vesicle (“yellow”), ribosome (“brown”), and mRNA splicing processes (“tan”). Finally, we were interested if the identified WGCNA modules were enriched for DEGs and DEPs suggesting their dynamic change in CUD. We found statistically significant enrichment of DEGs in all 4 CUD-associated RNA modules most prominently among downregulated DEGs (Figure 2D). Further, significant enrichment of DEPs was found in the three protein modules indicating differential network activity of RNA and protein co-expression modules in CUD (Figure 2E).

**Figure 2.**
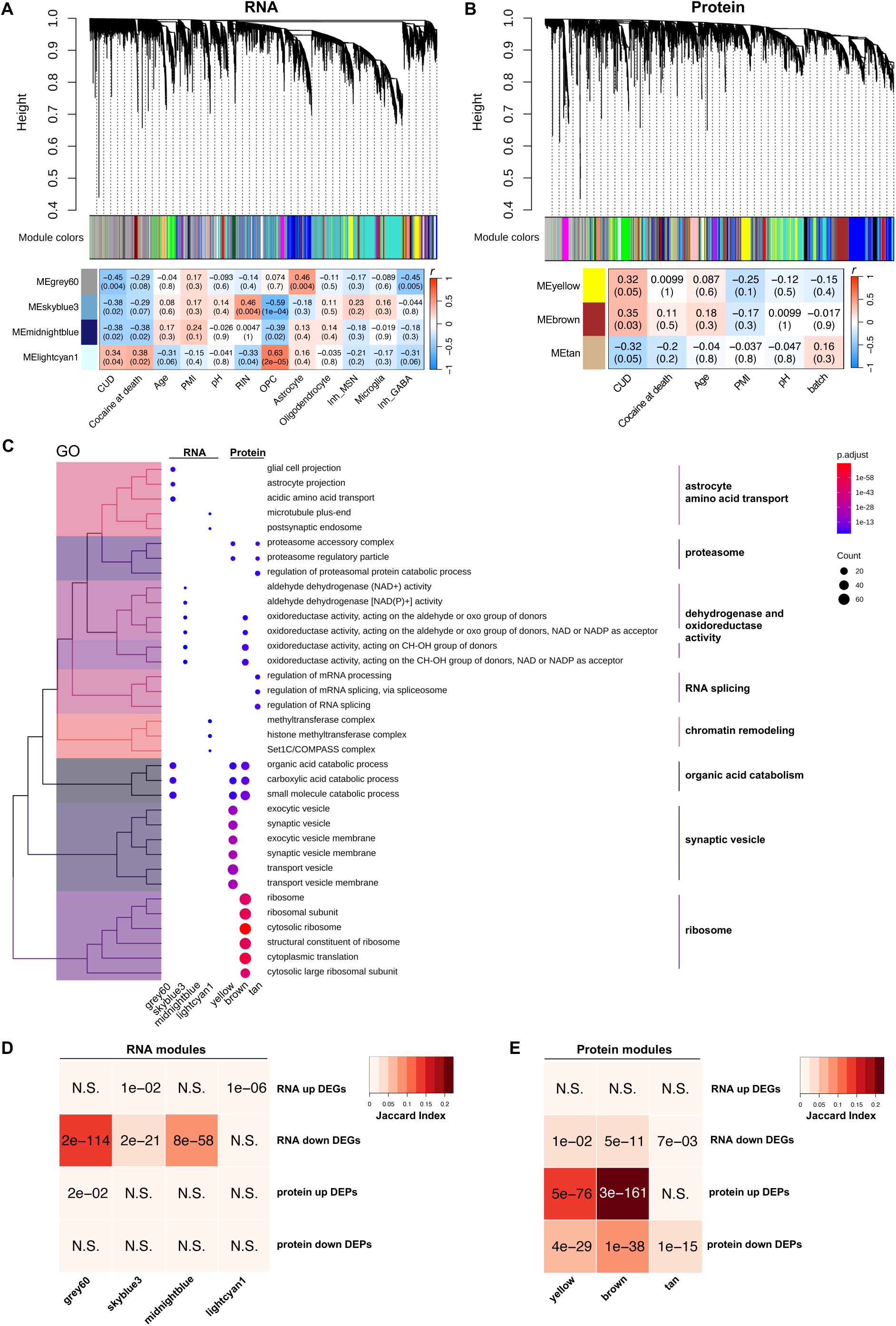
Identification and functional characterization of cocaine use disorder associated RNA and protein co-expression networks. Dendrogram resulting from weighted co-expression network analysis (WGCNA) in **A** RNA-seq and **B** proteomics datasets indicating assignment of co-expressed genes to modules with color labels. Module eigengene (ME) correlation with CUD, available covariates, and cell type estimates from deconvolution analysis for CUD-associated co-expression modules. Panels contain color-coded Pearson correlation coefficient *r*. Significance of correlation is shown in brackets. PMI: postmortem interval, pH: postmortem brain tissue pH value, RIN: RNA integrity number, OPC: oligodendrocyte progenitor cell, Inh_MSN: cell type estimates for DRD1- and DRD2-expressing medium spiny neurons (MSNs), Inh_GABA: cell type estimates for GABAergic, non-MSN interneurons, batch: proteomics processing batch. **C** functional characterization of co-expressed genes to identify shared and specific pathways among RNA and protein co-expression modules in CUD. A treeplot visualization of statistically significant results from GO enrichment analysis results of co-expression module genes is shown. Analysis of overlap between up- and downregulated DEGs and DEPs and co-expression modules for **D** RNA co-expression and **E** protein co-expression module genes. Panels contain p-values from a Fisher-Test indicating the significance of overlap. DEG: differentially expressed gene from RNA-seq (p_nominal_<0.05), DEP: differentially expressed protein (p_nominal_<0.05). N.S.: not significant.

### Factor analysis-based multi-omics integration of miRNA-seq, RNA-seq, and proteomic datasets confirms metabolic changes as a key hallmark of the CUD brain

As an additional multi-omics integration analysis of miRNA-seq, RNA-seq, and proteomic datasets, we performed Multi-Omics Factor Analysis (MOFA). Here, we aimed to identify a latent factor representation of our high-dimensional CUD dataset by an integrative “in parallel” analysis of the three omics datasets. MOFA inferred 10 latent factors (Figure 3A, Figure S4A). In total, the MOFA model explained 34%, 71%, and 61% of the variance in the miRNA-seq, RNA-seq, and proteomic datasets, respectively (Figure S4B). Correlation of factors with covariates revealed a significant association of factor 10 with CUD status (*r*=0.33, p=0.04, Figure 3B+C), a factor enriched for pathways involved in synaptic signaling and oxidative phosphorylation both having been reported as deregulated biological processes in human CUD^19,21,27,33^. The association with factor 10 was even stronger for cocaine at death (*r*=0.47, p=0.002, Figure S4C) suggesting factor 10 particularly reflects the intoxication state of CUD. Factor 10 was further associated with RNA-based cell type estimates for astrocytes (*r*=-0.38, p=0.02, Figure S4D), GABAergic inhibitory neurons (*r*=-0.39, p=0.01, Figure S4E) and oligodendrocytes (*r*=0.32, p=0.05, Figure S4F).

**Figure 3.**
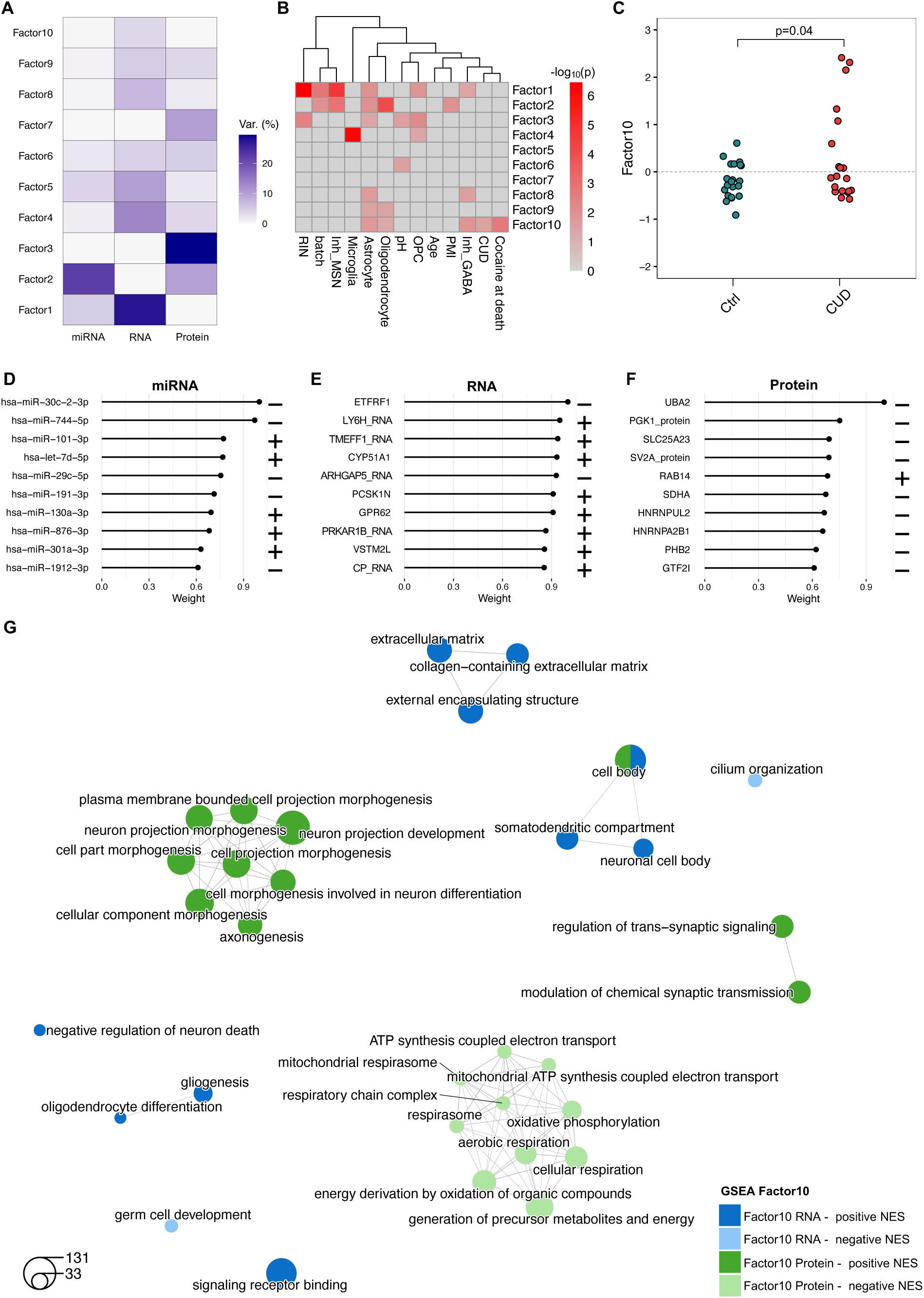
Integrative bulk-level analysis of miRNA-seq, RNA-seq, and proteomics data by multi-omics factor analysis reveals a CUD-associated latent factor enriched for synaptic and metabolic genes. **A** Variance explanation per factor for each omics dataset, miRNA-seq (all 1,542 miRNAs), RNA-seq (top 4,270 variable RNAs), and proteomics (all 4,270 proteins) based on the factor model identified in multi-omics factor analysis (MOFA). **B** Correlation of factors with CUD, covariates, and cell type estimates from cell type deconvolution analysis in RNA-seq data. Significance of correlation as -log_10_(p-value) is shown in the heatmap. **C** comparison of sample loadings on factor 10 identifies a significant difference between CUD cases and Ctrl individuals with CUD individuals having stronger positive loadings on this latent factor. p-value from a Wilcoxon-Test on factor 10 loadings is shown. Top 10 features i.e. **D** miRNAs, **E** RNAs, and **F** proteins with strongest absolute weights on CUD-associated factor 10. + indicates positive, - indicates negative weight. **G** gene set enrichment analysis (GSEA) results for positive and negative RNA (blue) and protein weights (green) on factor 10 identifies functional pathway modules among statistically significant GO terms (FDR q<0.05).

Next, we were interested in the biological processes represented by CUD-associated factor 10. For this, we inspected the weights of individual miRNAs, RNAs, and proteins on this factor (Table S8). hsa-miR-30c-2-3p and hsa-miR-101-3p were identified as the miRNAs with the strongest negative and positive weights, respectively (Figure 3D). At the RNA level, the top features were electron transport chain-associated gene *ETFRF1* for negative weights and nicotinic acetylcholine receptor modulator *LY6H* for positive weights (Figure 3E). At the protein level, UBA2 related to protein SUMOylation and the GTPase RAB14 involved in vesicle trafficking displayed the largest absolute negative and positive weights, respectively (Figure 3F). Using weight information from RNA and protein datasets, pre-ranked GSEA was performed to identify biological pathways associated with factor 10 (Table S9). Largest functional pathway modules after GO term clustering were related to neuronal morphology and oxidative phosphorylation (Figure 3G). Further, among significant results with positive GSEA normalized enrichment score (NES) at the protein level, synaptic signaling pathways emerged confirming results from the WGCNA analysis of the proteomic dataset. A second highly connected pathway cluster consisting of metabolic GO terms related to oxidative phosphorylation was specific to protein results with negative NES. Thus, while previous pathway enrichment analyses of DEGs, DEPs, and WGCNA module genes suggest alterations in fatty acid metabolism and oxidoreductase enzymes, MOFA identified additional alterations in oxidative phosphorylation depicting a shared downstream process of the previously identified metabolic processes.

### Single-nuclei RNA sequencing identifies the cell type-specificity of transcriptional changes and suggests astrocytes and medium spiny neurons as particularly deregulated cell types in CUD

Our bulk-level analyses in the VS provide evidence for metabolic, ribosomal, and synaptic changes in CUD. However, it remains unclear to what extent these findings are specific to or driven by individual cell types. To identify potential cell type-specific transcriptomic changes in CUD we performed snRNA sequencing in a subset of N=16 individuals (8 CUD cases, 8 control individuals, Table S1). In our dataset of N=20,759 single nuclei, we identified 12 cell type clusters in the VS by analyzing the expression levels of known marker genes (Figure 4A+B, Figure S5). Major neuronal cell types of the VS include DRD1 and DRD2-expressing medium spiny neurons (MSNs) as well as non-MSN GABAergic interneurons. Further, glial cell types such as astrocytes, oligodendrocytes, oligodendrocyte progenitor cells (OPC), and microglia were identified. To determine DEGs in the snRNA-seq dataset, we performed CUD vs. Ctrl differential expression analysis in each major cell type cluster identifying a total of 1,996 DEGs (|log2FC|>0.25, p_adj_<0.05, Table S10, Figure 4C). These include well-described immediate early genes such as *JUN* and *FOSB* that have been repeatedly described to be induced in human brain and rodent brain following cocaine exposure^34–36^. The strongest CUD-associated expression deregulation was found in D1-MSNs (674 DEGs), D2-MSNs (736 DEGs), and astrocytes (513 DEGs, Figure S6A) which is well reflected by cell type-specific DEG patterns in a rodent model of repeated cocaine intake^26^. As RNA expression data was available at the bulk- and single-cell level in our study, we compared CUD-associated DEG patterns between bulk RNA-seq and snRNA-seq (DE analysis across all clusters) using RRHO. Here, a strong convergence of results was observed (Figure S6B) confirming robust overlap between DEG signatures at bulk-level and cluster-ignorant snRNA-seq analysis. To investigate cell type-specific pathway deregulation, a GO enrichment analysis was performed using DEGs from each of the 7 major cell type clusters (Figure 4D, Table S11). Significant GO term clusters conserved across neuronal and glial cell types include “structural constituent of ribosome” (D1-MSN, D2-MSN, astrocyte, oligodendrocyte), oxidative phosphorylation pathways (D1-MSN, astrocyte), “cadherin binding” (D1-MSN, oligodendrocyte), and “ubiquitin protein ligase binding” (D1-MSN, OPC). Here, ribosomal and oxidative phosphorylation changes might be a conserved feature of CUD across striatal brain regions as they also emerged in a recent cell type-specfic analysis in the CN^27^. Further, we found significant pathway modules that were specific to either neurons or glial clusters such as immunological pathways related to T-cell receptor and MHC binding (microglia, OPC), calcium channel signaling (D1-MSN, D2-MSN), “3’-5’-cyclic-GMP-phosphodiesterase activity” (oligodendrocyte), and “glutamate receptor activity” (OPC).

**Figure 4.**
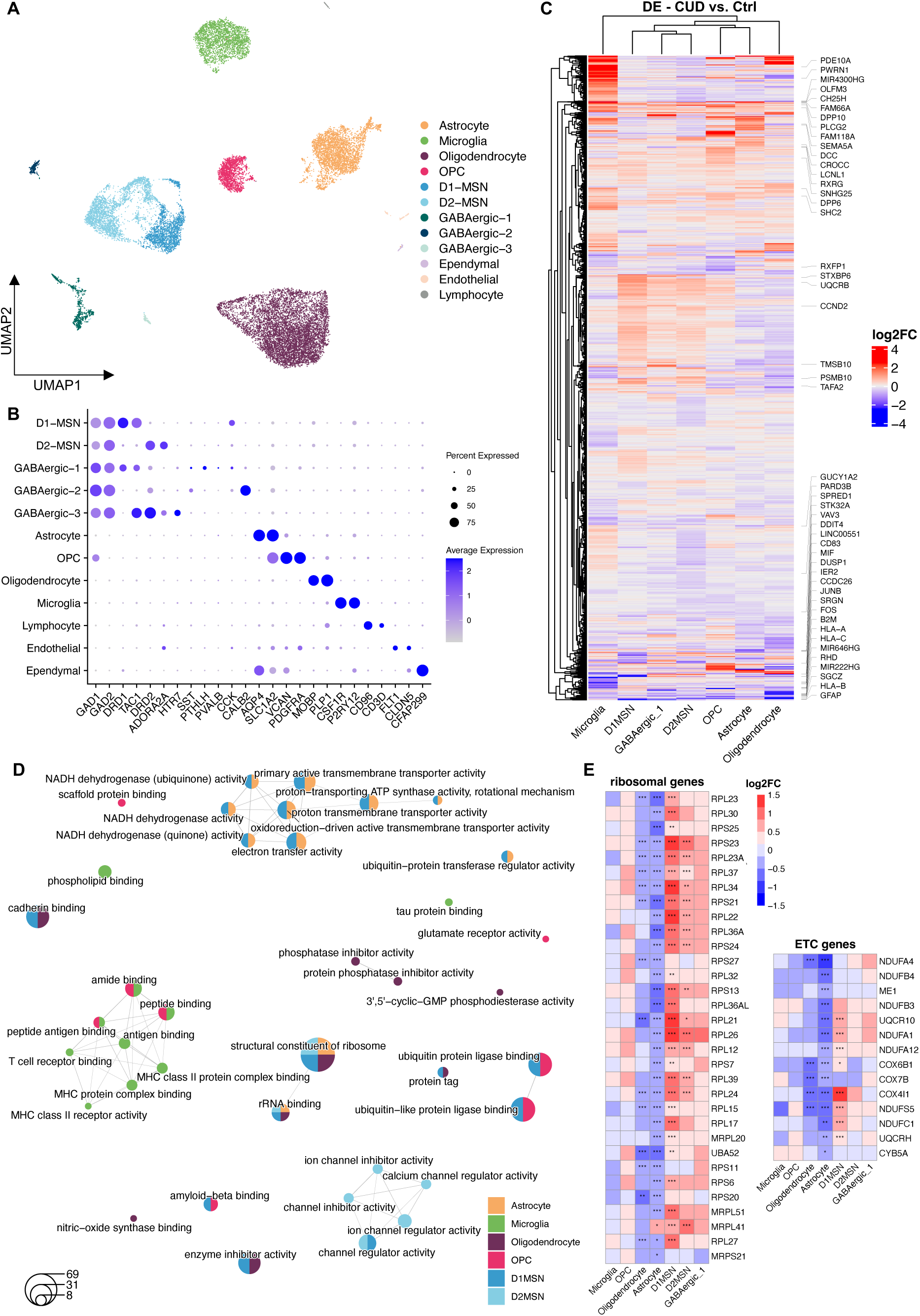
Single-nuclei RNA sequencing of the ventral striatum identifies cell type specific transcriptomic deregulation patterns in cocaine use disorder. **A** UMAP representation of the transcriptomic profiles of N=20,759 single nuclei from N=8 individuals with and N=18 individuals without CUD identifies 12 cell type clusters in the ventral striatum (VS). **B** annotation of cell types based on expression levels of known cell type marker genes. **C** heatmap of cell type specific differential expression (DE) in the 7 major cell types of the VS revealing N=1,996 DE genes (|log2FC|>0.25, FDR q<0.05). Top deregulated genes per cell type based on expression log2-fold change are labeled with gene name. **D** Statistically significant results from GO enrichment analysis (FDR q<0.05) of differentially expressed genes (DEGs) from major cell types indicate common and cluster-specific biological pathway deregulation. **E** heatmap of log2-fold change in the expression of pathway-defining genes for pathways characterized by highly significant enrichment of DEGs in neuronal and glial cell types. Ribosomal and electron transfer chain (ETC) genes were investigated. *p<0.05, **p<0.01, ***p<0.001.

As ribosomal and oxidative phosphorylation pathways were reported in existing literature^19,27^, have emerged as main findings in our bulk-level analyses, and were conserved across neuronal and non-neuronal clusters in snRNA-seq, we were interested in the direction of DEG patterns for these biological processes in individual cell types. For this, we evaluated log2FC and p-values of genes related to ribosomal and electron transport chain (ETC) pathways by extracting genes defining “structural constituent of ribosome” and “electron transfer activity” GO terms. For both, ribosomal and ETC genes, significant upregulation was detected in neurons, especially in D1-MSNs and D2-MSNs, while the same genes were significantly downregulated in glial cell types, most prominently in astrocytes and oligodendrocytes (Figure 4E). Cell type-specific analysis of gene expression in CUD thus suggests inverse deregulation patterns between neurons and glia cells within genes related to the ribosome and oxidative phosphorylation.

### Cell type-specific co-expression analysis identifies gene network alterations in astrocytes and striatal medium spiny neurons that are conserved across human CUD and a rodent model of repeated cocaine intake

Consistent with the analysis approach in bulk-level datasets, we aimed to identify and functionally characterize transcriptional co-expression networks in the single-nuclei dataset. We first performed a cell type-specific co-expression network analysis in our human snRNA-seq data using hdWGCNA^37^. Module prioritization (Table S12, STAR Methods) resulted in 8 cell type-specific co-expression modules characterized by significant downregulation in CUD (differential module eigengene log2FC<0, q<0.05, Figure S7A-D, Table S12). Cell type-specific co-expression modules were identified in astrocytes, MSNs (Inh_MSN), and non-MSN GABAergic interneurons (Inh_GABA, Table S13). Co-expression module genes were significantly enriched in cell type-specific DEGs, most prominently in downregulated DEGs, further supporting the downregulation of the identified cell type-specific co-expression networks in CUD (Figure S7E). Pathway enrichment analysis in astrocytic co-expression networks highlights deregulation of glutamatergic synapse and neurotrophic signaling as well as glutamate and fatty acid metabolism in CUD (“Astrocyte-M12”, “Astrocyte-M14”, Figure S8A+B, Table S14). In line with this, we found important regulators of glutamatergic signaling such as glutamate dehydrogenase *GLUD1* (“Astrocyte-M12”, Figure S7F), glutamate transporters *SLC1A2* and *SLC1A3* (“Astrocyte-M14”, Figure S7F) as well as metabotropic glutamate receptor *GRM3* (“Astrocyte-M14”, Figure S7F) among module hub genes in the astrocyte-specific CUD-associated co-expression modules. In neuron-specific co-expression modules, we identified metabolic pathways related to nucleoside and ketone body metabolism, ion transport processes, and GABAergic signaling (Figure S8C+D, Table S14). Module hub genes include several ionotropic neurotransmitter receptor genes such as GABA-A receptor subunits *GABRB1* and *GABRB3*, potassium channel gene *KCNQ5*, calcium channel subunit *CACNA1E* (all “Inh_MSN-M7”, Figure S7F), sodium-calcium exchanger *SLC8A1*, and *SLC22A17* involved in iron transport (both “Inh-MSN-M2”, Figure S7F).

Next, to investigate potential conservation patterns of network changes across human CUD and a repeated cocaine intake paradigm in rats, we performed consensus hdWGCNA in the N=20,759 human nuclei and N=11,288 nuclei of the ventral striatum (nucleus accumbens) from male Sprague-Dawley rats undergoing 7 days of cocaine exposure^20^ (Figure 5A). For MSNs and astrocytes, the cell types that have been most prominently implicated in human CUD based on DE and hdWGCNA analyses and also showed the largest number of DEGs in the rodent dataset, we found 7 (“Inh_MSN-CM1”, “Inh_MSN-CM4”, “Inh_MSN-CM5”, “Inh_MSN-CM6”, “Inh_MSN-CM7”, “Inh_MSN-CM9”, “Inh_MSN-CM13”) and 3 (“Astrocyte-CM4”, “Astrocyte-CM6”, “Astrocyte-CM8”) significantly CUD-associated consensus co-expression modules, respectively (Table S15+S16). In astrocytes, a strong overlap was observed between human module genes (“Astrocyte-M14”) and consensus module genes (“Astrocyte-CM4”, “Astrocyte-CM6”, Figure 5B+C). Consistent with the hdWGCNA analysis in human CUD, the 3 astrocyte-specific consensus modules displayed negative DME indicating downregulation of the consensus networks in CUD (Table S15). Astrocyte-specific consensus modules were enriched for pathways involved cAMP signaling, fatty acid metabolism, and glutamatergic signaling (Astrocyte-CM4”, “Astrocyte-CM6” Figure 5E+F, Table S17) supporting pathway results from human modules. In neurons, the strongest overlap was observed between human module “Inh_MSN-M7” and consensus module “Inh_MSN-CM4” that was also downregulated in CUD (DME log2FC=-0.20, Figure 5B+D, Table S15). At the pathway level, nucleoside and fatty acid metabolism changes were among the significant findings in neuron-specific consensus module “Inh_MSN-CM4” (Figure 5G+H, Table S17) in line with results from the human module “Inh_MSN-M7”. We further detected pathway enrichment related to synaptic vesicle transport and long-term depression in “Inh_MSN-CM4” indicating conserved deregulation of neuronal functions. Notably, KEGG enrichment analysis of consensus co-expression module genes revealed multiple SUD-related pathways among the most significant associations such as “Morphine addiction”, “Nicotine addiction”, and “Cocaine addiction” in both, astrocytes and MSNs, confirming the presence of addiction-relevant genes in consensus co-expression modules (Figure 5E-H). Results from the consensus network analysis thus indicate a set of deregulated biological processes including fatty acid metabolism and glutamatergic signaling conserved between human CUD and a rat model of repeated cocaine intake.

**Figure 5.**
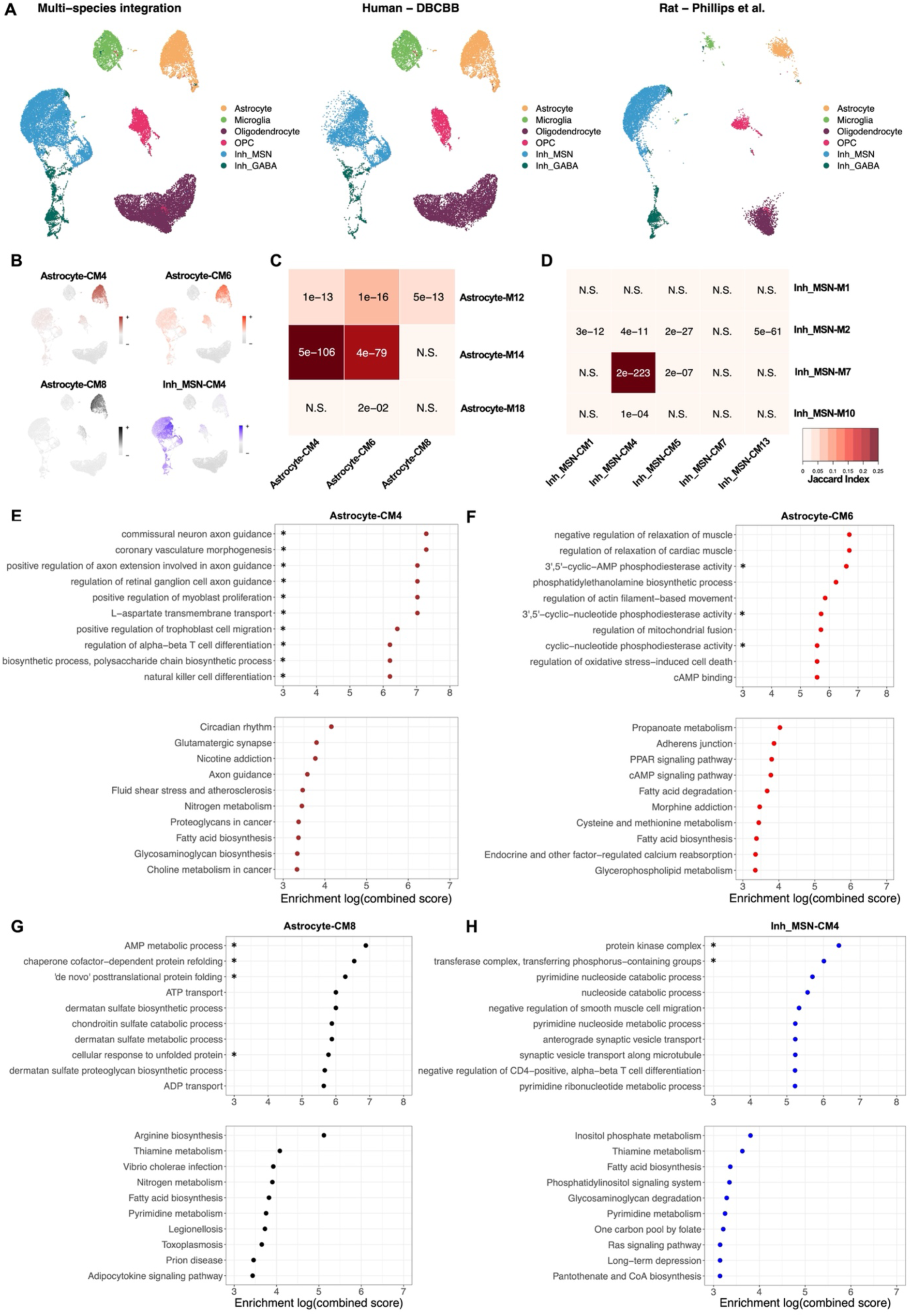
Integrative analysis of snRNA-seq datasets from human cocaine use disorder and a rat model of repeated cocaine intake reveals consensus co-expression modules across species. **A** UMAP representations of the integrated snRNA-seq dataset containing N=20,492 human and N=9,649 rat nuclei of ventral striatum cell major types. DBCBB – Douglas Bell Canada Brain Bank, rat dataset from Phillips et al. 2023, Mol Cell Neurosci. **B** expression of module eigengenes from consensus co-expression modules with significant DME in CUD and cell type specific expression. Analysis of the relationship between human-only and consensus co-expression modules in **C** astrocyte and **D** Inh_MSN clusters measuring the overlap of co-expression module genes. Jaccard index and p-values from Fisher-Test are shown. N.S.: not significant. GO and KEGG enrichment analysis results for consensus co-expression modules **E** Astrocyte-CM4, **F** Astrocyte-CM6, **G** Astrocyte-CM8, and **H** Inh_MSN-CM4 characterized by strongest overlap with modules identified in human CUD. Combined scores for results ranking were calculated from enrichment statistics in EnrichR, *FDR q<0.05.

### Altered astrocyte-neuron crosstalk in human CUD affects glutamatergic signaling and communication via cell adhesion molecules

Most prominent changes in CUD-associated gene expression and co-expression networks were found in astrocytes and MSNs of the VS. To better understand the consequences of transcriptional deregulation in these cell types, we were interested in their molecular crosstalk. For this, CellChat was performed in astrocyte, D1-MSN, and D2-MSN clusters to identify the differential activity of ligand-receptor (LR) interaction pairs in CUD. We found upregulation of 16 and downregulation of 106 LR pairs in CUD (Table S18).

Upregulated signaling in CUD was mainly related to secreted signaling factors such as glycoprotein *SLIT2* or Eph receptor ligand *EFNA5*, originating from D1-MSNs (Figure 6A). D1-MSNs and D2-MSNS were also the main receiving cell type of upregulated signaling in CUD indicating limited neuron-astrocyte crosstalk through upregulated LR pairs. In contrast, neuron-astrocyte crosstalk was particularly present among downregulated LR pairs with the strongest expression changes of ligands detected in astrocytes and D2-MSNs (Figure 6B). The astrocytic ligand expression profile was characterized by the downregulation of glutamatergic signaling genes (*SLC1A2, SLC1A3*) in line with results from astrocyte-specific co-expression modules and supported by consistent findings of *SLC1A2* downregulation in rodent models of cocaine addiction^38^. Further, among downregulated ligands on astrocytes, we found several cell-cell adhesion molecules including *NRXN1* and *CADM1* that have an important role in securing the structural integrity of tripartite synapses^39^. Both, D1-MSNs and D2-MSNs were receivers of altered glutamate signaling from astrocytes as subunits of AMPA (*GRIA3*, *GRIA4*) and kainate receptors (*GRIK2*, *GRIK3*) as well as metabotropic glutamate receptor *GRM7* were downregulated in MSNs. Further, D2-MNSs were the main receivers of differential cell-cell adhesion signaling as they expressed reduced levels of NRXN interaction partners *NLGN1* and LRRTM family members *LRRTM3* and *LRRTM4*. Thus, our analysis of differential LR pair expression in CUD suggests altered astrocyte-neuron crosstalk related to the downregulation of glutamatergic and cell adhesion signaling. Interestingly, the same set of glutamatergic and cell adhesion signaling genes have been identified as module hub genes of hdWGCNA co-expression modules “Astrocyte-M14” and “Inh_MSN-M7” (Figure S7F) that were both downregulated in CUD and displayed conservation patterns across species. CellChat analysis thus confirms findings from co-expression analysis and further outlines altered crosstalk between astrocytes and MSNs as an important hallmark of the VS in CUD.

**Figure 6.**
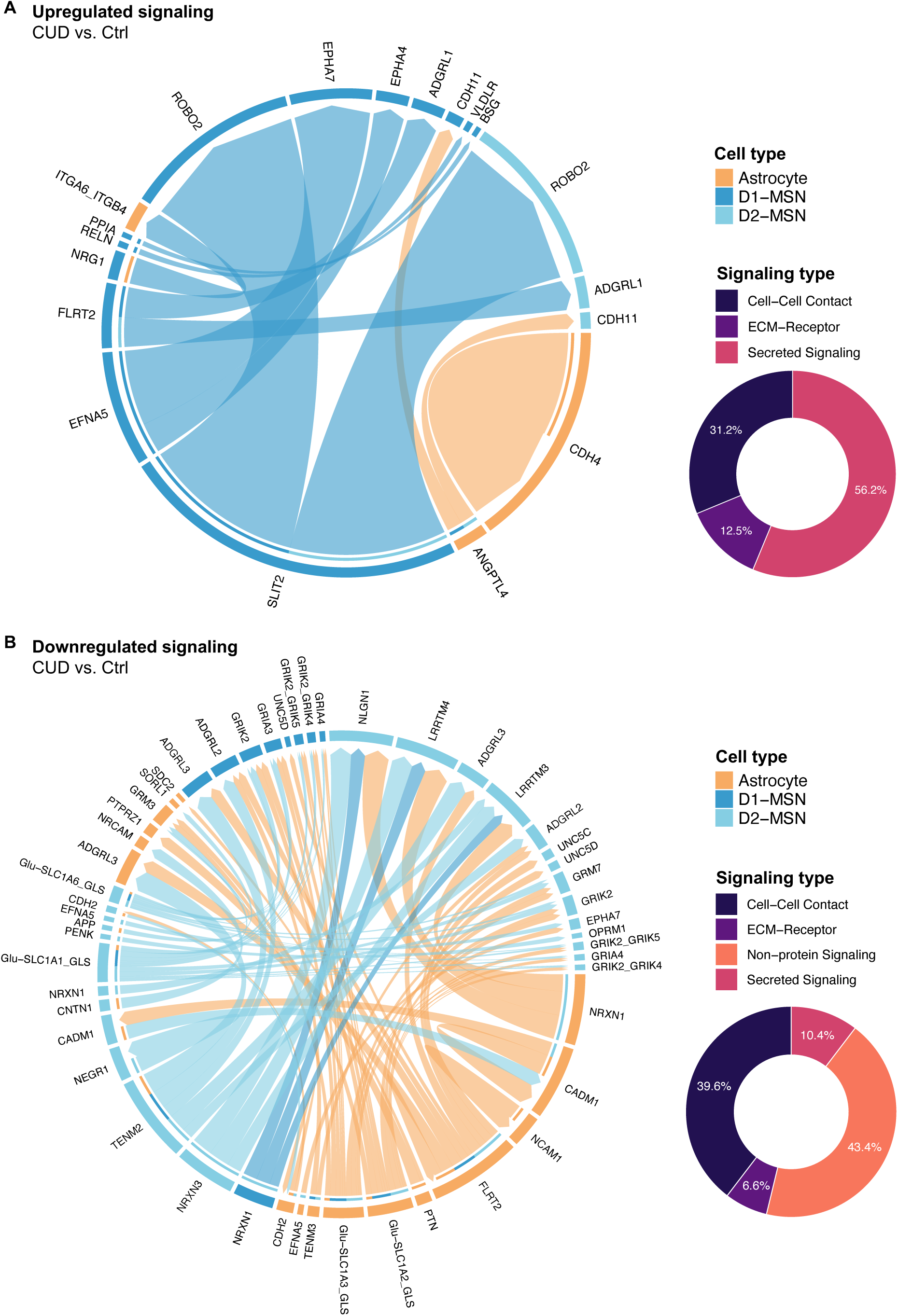
Astrocyte-neuron crosstalk in cocaine use disorder is characterized by aberrant glutamatergic and cell-cell adhesion signaling. Astrocytes and medium spiny neurons (MSNs) displayed the strongest CUD-associated changes in differential gene expression and cell type specific co-expression module activity. Differential expression analysis of ligand-receptor pairs was performed using CellChat to evaluate differential astrocyte-neuron crosstalk patterns in CUD. **A** circos plot for significantly upregulated ligand-receptor pairs in astrocytes (orange), D1-MSNs (blue) and D2-MSNs (light-blue). Arrows indicate the directionality of signaling from a ligand expressed in the sender cell to its receptor expressed in the receiver cell type (arrowhead). Percentage of signaling types among upregulated ligand-receptor pairs are provided in the donut plot. **B** circos plot for significantly downregulated ligand-receptor pairs in CUD for astrocytes (orange), D1-MSNs (blue) and D2-MSNs (light-blue). Also, the distribution of signaling types within downregulated crosstalk was analyzed and visualized using a donut plot. Cut-offs for significant differential expression of ligand-receptor pairs were a minimum of 5% change in ligand expression (|log2FC>0.07|) and receptor differential expression into the same direction, both at 5% FDR significance.

## Discussion

The present study depicts an integrative multi-omics analysis framework for characterizing neurobiological changes in the human ventral striatum in CUD. To our knowledge, our study is the first in a psychiatric phenotype that performed multi-omics integration of microRNA-seq, RNA-seq, and proteomics datasets together with cell type-specific analyses by snRNAseq in postmortem human brain tissue. Our unique study design enables the identification of molecular changes in CUD at different levels of biological regulation – starting from miRNAs that act on the RNA level up to the protein level. This allowed us to provide novel insights into across-omics conservation patterns of molecular deregulation in CUD, for instance, involving fatty acid metabolism. By performing additional analyses at single-cell resolution, we showed that bulk-level results, such as ribosomal and oxidative phosphorylation changes, were also reflected in the single-nuclei dataset. At the same time, we gained important insights into directional effects of transcript deregulation patterns in glial compared to neuronal cell types, highlighting the value of cell type-specific analyses. Finally, we show the advantage of integrating rodent model and human datasets where we found converging evidence for altered glutamatergic signaling in CUD that was also confirmed by an astrocyte-neuron crosstalk analysis.

Pathway associations with CUD from bulk-level analyses include synaptic, immunological, ribosomal, and metabolic changes related to fatty acid metabolism and oxidative phosphorylation. This confirms findings from previous analyses of miRNA, RNA transcriptomic, and proteomic studies in CUD^15,17,19–22,27,33^. Importantly, our study provides the first insights into the inter-relationship between layers of biological regulation in CUD by identifying across-omics conservation, especially of the fatty acid metabolism changes as indicated by DEG/DEP overlap and WGCNA co-expression networks. This underpins the deregulation of fatty acid metabolism as a central metabolic feature of the CUD brain and suggests that these metabolic alterations previously observed at the RNA level^30,33^ also extend to the protein level. While convergent evidence across omics analyses was observed, we also found several biological processes exclusively detected in single-omics analyses. From this observation, we conclude that integrative multi-omics analyses depict an important approach for a better understanding of molecular changes in complex phenotypes such as CUD, as single-omics analysis might miss important disease associations.

A frequent criticism of gene expression studies is that transcript levels are poorly predictive for protein levels thus limiting the interpretability of findings. Our transcriptome-proteome correlation analysis revealed an overall moderate correlation (*r*=0.43) between RNA and protein levels in the VS which is well in line with results from a study investigating schizophrenia-associated molecular changes in the prefrontal cortex that also observed a correlation of *r*=0.43 between transcriptome and proteome^32^. Importantly, we found the strongest positive correlation between RNA and protein levels for genes involved in neuronal function and synaptic signaling such as *PDE10A*, *SCN4B*, and *FKBP5*, suggesting that the assumption of RNA levels predicting protein levels holds true for these frequently studies genes in psychiatric disorders. Strong negative RNA-protein correlation was detected for genes involved in oxidative phosphorylation. While this might be a true biological effect, it could also be a consequence of insufficient mitochondrial protein solubilization during brain tissue lysis.

In cell type-specific DEG analyses, we found inverse deregulation patterns between neuronal and glial cell types related to ribosomal and oxidative phosphorylation pathways. Alterations of these pathways in CUD is in line with results from previous postmortem human brain studies^19,27^. In a cell type-specific multi-omics study of CUD using snRNA-seq and snATAC-seq, we found similar ribosomal gene deregulation patterns in the caudate nucleus^27^. One explanation for a neuron-specific upregulation of ribosomal genes in CUD could be an increased demand for local translation particularly in neurons as ribosomes are abundant in dendrites and synapses where they are required for the local synthesis of proteins involved in neurotransmission and synapse structure^40^. Our finding of increased ribosomal protein levels based on bulk-level proteomics in the VS further supports the hypothesis of an increased number of ribosomes in CUD. Further, differential expression of oxidative phosphorylation genes was detected in a bulk-level RNA-seq study of the hippocampus in CUD^19^. While the authors report on the overall downregulation of oxidative phosphorylation genes, we show that this finding is cell type dependent in the VS with downregulation patterns observed for glial cell types while the same genes were upregulated in neurons. We further found fatty acid metabolism changes to be especially prominent among astrocyte-specific co-expression modules highlighting the value of cell type-specific analyses to better understand findings that have been previously reported in bulk-level analyses. Follow-up studies are required to investigate if ribosomal and oxidative phosphorylation changes depict direct effects of cocaine exposure or are compensatory effects induced by long-term cocaine intake.

Interestingly, we did not observe neurotransmission or synaptic plasticity pathways in the differential transcript expression analysis of bulk- or snRNA-seq datasets of the VS. In contrast, bulk-level RNA-seq analysis of the prefrontal cortex subregion Brodmann Area 9 (BA9) in a subset of N=25 individuals from our DBCBB cohort revealed strong transcriptional changes for synaptic signaling genes in CUD^33^. Also, in the snRNA-seq/snATAC-seq study of CUD in the caudate nucleus, a brain region involved in compulsive drug intake patterns harboring similar cell types as the VS, we found consistent alterations in synaptic and ion channel signaling across transcriptomic and chromatin accessibility datasets^27^. At the same time, metabolic findings related to fatty acid metabolism or oxidative phosphorylation were consistently associated with CUD across brain regions. We hypothesize that this observation could reflect an aspect of late-stage CUD where the ventral striatum has undergone profound neuroadaptations in response to chronic cocaine intake but resides in an anaplastic state characterized by suppression of long-term depression (LTD)^41^. This is also supported by the MSN-specific co-expression analysis where vesicle transport and LTD pathways emerged for the CUD-downregulated consensus module “Inh_MSN-CM4”.

A strong advantage of rodent models in addiction research is controlled experimental conditions, allowing mechanistic studies on drug effects by systematic exclusion of confounding factors. To complement the study of postmortem brains, which is associated with strong interindividual heterogeneity, with a controlled laboratory experiment, we performed integration of our human snRNA-seq data with snRNA-seq from a rat model of repeated cocaine intake. Here, we deciphered conserved CUD-associated gene networks in the VS. Most prominent network findings were glutamatergic signaling alterations that were conserved across human and rat datasets as they emerged in consensus hdWGCNA and astrocyte-neuron crosstalk analysis. Craving and relapse, two key symptoms of CUD, have been shown to correlate with aberrant glutamate signaling in the VS^42^ and our study identified astrocytes and MSNs as important cell types that may contribute to this observation. While D1- and D2-MSNs have been recognized early on as important cell types for addiction research, there is increasing evidence for a key role of astrocytes in CUD^43,44^. For instance, the neurovascular effects of cocaine characterized by cerebral blood flow reduction were shown to be dependent on astrocytic calcium signaling and chemogenetic inhibition of astrocytes prevented cerebral vasoconstriction following a cocaine challenge in rodents^45^. Regarding glutamatergic signaling, glutamate transporter GLT-1 (SLC1A2) was consistently downregulated in NAc astrocytes of rats repeatedly exposed to cocaine and experimental normalization of GLT-1 levels reduced reinstatement of cocaine seeking^38,46,47^. Our study confirms a downregulation of *GLT-1* in in human CUD astrocytes while also suggesting ionotropic and metabotropic glutamate receptor subunits to be downregulated in D1- and D2-MSNs. Another important finding of the altered astrocyte-neuron crosstalk in CUD was changes in cell-cell adhesion dynamics that could further enhance abnormal glutamatergic signaling, as reduced NRXN-NLGN interaction disrupts the structural integrity of tripartite synapses, thereby additionally impeding glutamate homeostasis^48,49^. Furthermore, the glutamatergic imbalance in the VS depicts a possible link to the oxidative phosphorylation changes for which we observed different directional effects in glia and neurons. For instance, conversion of glutamate to α-ketoglutarate was shown to fuel the tricarboxylic acid cycle in brain mitochondria thereby temporarily increasing ATP production via oxidative phosphorylation changes^50^. Further studies are required to disentangle the direct and adaptive effects of aberrant glutamatergic signaling with the aim to investigate the modulation of glutamate system genes as a potential pharmacotherapy in CUD.

While aiming for the largest sample size possible, a limitation of the present study is the relatively small discovery cohort that might not represent the full spectrum of the CUD phenotype. Strong inter-individual heterogeneity depicts a general phenomenon in CUD, as cocaine is frequently used together with other drugs of abuse such as alcohol, cannabis, or opioids^51^. While our cohort was selected to be homogenous in sex and ancestry, it cannot be excluded that exposure to other drugs of abuse or medication might interfere with CUD-associated differential expression signatures. With the available sample size, we were not able to adjust for all potential influences on miRNA, RNA, and protein expression levels but additional covariates could be included in the analysis of larger cohorts. Further, for the interpretation of findings, it needs to be considered that the cross-sectional design does not allow to distinguish between cumulative effects of cocaine exposure and compensatory neuroadaptations.

Future studies should focus on the identification and validation of disease mechanisms based on multi-omics analyses for instance by identifying master regulators of CUD-associated transcriptional changes in individual cell types^27^. Further, multi-omics integration and conservation analysis between human CUD and rodent models capturing addiction-like criteria should be performed to characterize conserved patterns of molecular deregulation thereby addressing the inherent limitations of the two study types^52^. In summary, such multi-omics-to-mechanism studies provide a powerful analytical framework for the identification of disease mechanisms and potential new therapeutic targets in CUD.

## Methods

### Human postmortem brain tissue

Postmortem human brain tissue of the ventral striatum from N=20 individuals with cocaine use disorder and N=21 unaffected individuals was obtained from the Douglas Bell Canada Brain Bank (DBCBB) in Montreal, Canada. Tissue sampling and acquisition of phenotype information at DBCBB was performed based on their established ethical guidelines. Our multi-omics study was approved by the Ethics Committee II of the University of Heidelberg, Medical Faculty Mannheim, Germany, under the register number 2021-681. Inclusion criteria were donor age > 18 and a DSM-IV diagnosis of cocaine dependence. Instead of cocaine dependence, throughout this study, the more recent terminology from DSM-V i.e. cocaine use disorder is used. Exclusion criteria were other substance use disorders except alcohol use disorder and a diagnosis of severe neurodevelopmental or neuropsychiatric disorders except major depressive disorder (MDD). To address strong inter-individual heterogeneity of human individuals and its effects on statistical analyses, we aimed for a homogenous sample in our study resulting in a final cohort of N=41 male individuals with European American ancestry consisting of N=20 individuals with CUD and N=21 non-affected individuals assigned to the Ctrl group. Phenotype information other than CUD status include tissue donor age, sex (reported as biological sex), ethnic background, postmortem interval (PMI), brain pH value, additional psychiatric diagnoses, toxicology at death and cause of death. A detailed description of phenotypes is provided in Table S1. Prior to all postmortem brain sample processing steps during omics data generation, randomization based on CUD status was performed to minimize confounding technical and batch effects.

### miRNA sequencing

Extraction of miRNAs from was performed using the miRNeasy Tissue/Cells Advanced Micro Kit (Qiagen, Hilden, Germany) resulting in a total RNA preparation that contains small RNAs such as miRNAs as well as long RNAs. For all postmortem brain samples, a total of 5mg tissue was used during total RNA extraction. RNA quality of the N=41 total RNA preparations was assessed on an Agilent TapeStation 4200 (Agilent, Santa Clara, CA, USA) and N=40 samples remained based on an RNA integrity number cut-off of > 5.5. Library preparation for miRNA-sequencing was performed using the NEXFLEX small RNA-seq kit v3 (Perkin Elmer, Waltham, MA, USA). Small-RNA libraries were sequenced on an Illumina NovaSeq 6000 (Illumina, San Diego, CA, US) with an average of 10 million reads per sample (1×50bp).

### RNA sequencing

RNA was extracted as total RNA from 5mg of postmortem human brain tissue using the miRNeasy Tissue/Cells Advanced Micro Kit (Qiagen, Hilden, Germany). RIN values were determined as previously described in the miRNA extraction procedure with RIN>5.5 as the RNA quality cut-off resulting in N=40 samples for which library preparation was performed. Transcriptomic profiling of the VS samples was based on an NEBNext Ultra II Directional RNA Library Prep Kit (New England Biolabs, Ipswich, MA, USA) that was used for library preparation after rRNA depletion. RNA sequencing for the N=40 samples was performed on an Illumina NovaSeq 6000 device with an average of 60 million read pairs (2×100bp).

### Single-nuclei RNA sequencing

Nuclei were isolated from N=8 CUD cases and N=8 unaffected Ctrl individuals. Samples for snRNA-seq were selected from the full cohort of N=41 individuals by the amount of available tissue to perform all preparations (miRNA, RNA, protein, and snRNA) from the same tissue sample. Samples with low postmortem interval and higher RIN value as measured in the bulk RNA-seq analysis were preferred. For the nuclei isolation, 10mg of frozen postmortem tissue was used to minimize the amount of free-floating debris. Samples were processed as indicated in the 10X Genomics Chromium Nuclei Isolation kit with RNase Inhibitor manufacturers protocol (10x Genomics, Pleasanton, CA, USA). Briefly, the tissue was dissociated in lysis buffer, filtered, and the cellular debris were removed. After multiple centrifugation and washing steps, a clean nuclei suspension was obtained. Nuclei were automatically counted on LUNA-FL™ Dual Fluorescence Cell Counter while excluding particles smaller than 5µm and bigger than 15µm of diameter from the counting. Single-nuclei RNA-seq (snRNA-seq) libraries were generated using the 10X Genomics Chromium Next GEM Single Cell 3ʹ Kit v3.1 (Dual Index). A total of 10,000 nuclei per sample were loaded into chromium chips. Libraries were prepared following the provided 10X Genomics Dual Index Kit protocol and sequenced using an Illumina NovaSeq6000 device with 50,000 read pairs per cell.

### TMT quantitative proteomics

Proteomic profiling was based on a Radioimmunoprecipitation Assay (RIPA) buffer lysate of postmortem human brain tissue samples. For each sample preparation, 50-70mg postmortem brain tissue was mechanically disrupted and homogenized under a liquid nitrogen atmosphere. Disrupted tissue was transferred to a reaction tube and was suspended in lysis buffer Pierce^®^ RIPA (Thermo Fisher Scientific, Waltham, MA, USA) with protease inhibitor (cOmplete^TM^ Mini, Roche, Basel, Switzerland). After a lysis period of 30min at 4°C, the suspension was centrifuged for 10 min at 12,500xg (4°C) and the supernatant was kept for further lysate preparation steps. Protein concentration was measured for each sample on Direct Detect^®^ Assay-free membrane cards using a Direct Detect^®^ Spectrometer (both Merck Millipore, Burlington, MA, USA). RIPA lysate containing 50µg total protein was processed for SDS gel electrophoresis. NuPAGE™ 4x LDS Sample Buffer and NuPAGE™ 10x Reducing Agent (both Thermo Fisher Scientific, Waltham, MA, USA) were added to the protein lysate and incubated for 10 min at 75°C. From this sample preparation, a total of 25µg protein were analyzed in TMT proteomics.

The reduction of disulfide bonds on cysteine was conducted using dithiothreitol (56°C, 30 minutes, 10 mM in 50 mM HEPES, pH 8.5) followed by alkylation with 2-chloroacetamide (room temperature (RT), in the dark, 30 minutes, 20 mM in 50 mM HEPES, pH 8.5). The SP3 protocol ^53,54^ was employed for sample clean-up, and trypsin (sequencing grade, Promega, Madison, WI, USA) was added at an enzyme-to-protein ratio of 1:50 for overnight digestion at 37°C (in 50 mM HEPES). The N=40 samples were randomized into three TMT multiplex batches (two with 16 samples each and one with 10 samples) based on CUD status and phenotypic covariates. Peptides were labelled with either TMT10plex (N=8 sample batch)^55^ or TMT16plex (2x N=16 sample batches)^56^ Isobaric Label Reagent (Thermo Fisher Scientific, Waltham, MA, USA) according the manufacturer’s instructions. In brief, 0.8 mg of reagent was dissolved in 42 ml of acetonitrile (100%) and 8 µl was added, followed by an incubation period of 1 hour at room temperature. The samples were incubated with 8 µl of 5% hydroxylamine for 15 minutes at room temperature. The samples from a given measurement batch were combined and desalted on an OASIS® HLB µElution Plate (Waters, Milford, MA, USA). The offline high pH reverse phase fractionation was conducted on an Agilent 1200 Infinity high-performance liquid chromatography system, equipped with a Gemini C18 column (3 μm, 110 Å, 100 x A 1.0 mm Phenomenex Torrance, CA, USA column was installed with a Gemini C18 4 x 2.0 mm SecurityGuard cartridge (Phenomenex, Torrance, CA, USA) as a guard column. The binary solvent system comprised 20 mM ammonium formate (pH 10.0) (A) and 100% acetonitrile as the mobile phase (B). The flow rate was set to 0.1 mL/min. Peptides were separated using a gradient of 100% A for 2 min, to 35% B in 59 min, to 85% B in another 1 min and kept at 85% B for an additional 15 min, before returning to 100% A and re-equilibration for 13 min. A total of 48 fractions were collected which were subsequently pooled into 12 fractions. Pooled fractions were dried under vacuum centrifugation, reconstituted in 10 μL 1% formic acid, 4% acetonitrile and then stored at −80 °C until LC-MS analysis.

An UltiMate 3000 RSLC nano LC system (Dionex, Sunnyvale, CA, USA) fitted with a trapping cartridge (µ-Precolumn C18 PepMap 100, 5µm, 300 µm i.d. x 5 mm, 100 Å) and an analytical column (nanoEase™ M/Z HSS T3 column 75 µm x 250 mm C18, 1.8 µm, 100 Å, Waters, Milford, MA, USA) was coupled to an Orbitrap Fusion™ Lumos™ Tribrid™ Mass Spectrometer (Thermo Fisher Scientific, Waltham, MA, USA) using the Nanospray Flex™ ion source in positive ion mode. The samples were applied to the trapping column at a constant flow rate of 30µL/min with 0.05% trifluoroacetic acid in water for a period of four minutes. After switching in line with the analytical column peptides were eluted at a constant flow of 0.3 µL/min using the method described below. The binary solvent system comprised 0.1% formic acid in water with 3% DMSO (solvent A) and 0.1% formic acid in acetonitrile with 3% DMSO (solvent B). The percentage of solvent B was increased from 2% to 8% in 4 min (N=10 TMT batch) or in 2 min (N=16 TMT batches), from 8% to 28% in 72 min (N=10 TMT batch) or 104 min (N=16 TMT batches), to 38% (N=10 TMT batch) or to 40% (N=16 TMT batches) in another 4 min and finally to 80% B in 4 min, followed by re-equilibration back to 2% B in 4 min. The peptides were introduced into the mass spectrometer via a Pico-Tip Emitter 360 µm OD x 20 µm ID; 10 µm tip (New Objective, Woburn, MA, USA) and an applied spray voltage of 2.4 kV. The capillary temperature was set to 275°C. A full mass scan was conducted with a mass range of 375-1500 m/z in profile mode in the orbitrap with a resolution of 120000. The maximum filling time was set at 50 ms, with a limit of 4×10^5^ ions. The data-dependent acquisition (DDA) was conducted with the Orbitrap resolution set to 30000, with a fill time of 94 ms, and a limitation of 1×10^5^ ions. A normalized collision energy of 36 (N=10 TMT batch) or 34 (N=16 TMT batches) was applied. MS^2^ data was acquired in profile mode. The define first mass was set to 110 m/z.

### Statistical analysis

All statistical analyses were performed in the statistical computing environment R, version 4.2.1. If not otherwise stated, Benjamini-Hochberg (FDR)^57^ correction was performed to adjust for multiple testing.

### Sequencing data preprocessing

Raw sequencing data from miRNA-sequencing were processed using FastQC v.0.12.1 and quality metrics were inspected. Sequencing adapters were trimmed using TrimGalore v.0.6.10. with automatic adapter detection and a minimum read length threshold of 18bp. Success of adapter trimming was evaluated in a post-processing run of FastQC. Alignment of miRNA-sequencing reads to the human reference genome (hg38) was performed using STAR v. 2.7.10b^58^. First, a genome index for the GENCODE GRCh38 primary assembly reference genome together with the GENCODE v.43 genome annotation file was created using STAR with --sjdbOverhang 49 for miRNA-sequencing data. Next, alignment was performed for the N=40 fastq-files while filtering for multi-mapping using the -- outFilterMultimapNmax 20 flag. Quantification of miRNAs was performed using featureCounts as implemented in Rsubread v.2.12.3^59^ with the miRNA-specific hsa.gff3 reference file from miRBase (https://www.mirbase.org/download/, release 22.1).

For the RNA-seq dataset, raw fastq-files were inspected in FastQC as described for the miRNA-sequencing dataset. The GRCh38 reference genome was indexed in STAR using the –sjdbOverhang 100 flag. Alignment in STAR was performed using default parameters. For the quantification of transcripts, featureCounts was used with the GENCODE v.43 genome gtf-file as the reference annotation.

Raw snRNA-seq data from the N=16 VS tissue samples was processed using Cell Ranger v.7.1.0 (10x Genomics)^60^ and resulting feature-barcode were imported in Seurat v.5.0.1^61^. Low quality nuclei were removed from the analysis using the following QC parameters: 900 < nFeature_RNA < 8500 and mitochondrial gene fraction <10%. After QC, ambient RNA correction was performed using SoupX v.1.6.2^62^. Expression data matrices from the N=16 samples were merged into a single Seurat object. Count normalization was performed using “NormalizeData”, followed by the identification of variable features (“FindVariableFeatures”) and data scaling (“ScaleData”). Next, “RunPCA” and “RunUMAP” were applied for dimensionality reduction and the N=16 individual datasets were integrated using “IntegrateLayers” with “method = HarmonyIntegration”. “FindClusters” was applied with a resolution parameter of 0.1. The association of clusters with technical parameters such as number of features and mitochondrial gene counts was inspected leading to the removal of one cluster that was strongly enriched for mitochondrial genes. Re-clustering at a resolution of 0.12 resulted in the final object with N=12 cell type clusters based on expression data from N=20,759 nuclei.

Proteomics data preprocessing and analysis was performed based on an adapted version of an analysis pipeline by Frank Stein (EMBL, Heidelberg, Germany). Preprocessing of the proteome data was performed using Fragpipe v20.0 (MSFragger v.3.8)^63^ by searching against a *Homo sapiens* proteome database (UP000005640, October 2022, 20594 entries) plus common contaminants and reversed sequences. The following modifications were included into the search parameters: Carbamidomethyl on Cysteine and TMT10/16 on lysine as fixed modifications, protein N-term acetylation, oxidation on methionine and TMT10/16 on N-termini as variable modifications. A mass error tolerance of 20 ppm was applied to both precursor and fragment ions. Trypsin was set as protease with a maximum of two missed cleavages. The minimum peptide length was set to seven amino acids. At least two unique peptides were required for protein identification. The false discovery rate on peptide and protein level was set to 0.01. The raw reporter intensities from the three TMT plexes were extracted from the raw tsv output files of FragPipe. Contaminants were removed, and only proteins quantified with at least two Razor peptides were included in the analyses. Due to the intrinsic nature of postmortem human brain studies characterized by strong interindividual heterogeneity, we did not perform imputation of missing values but restricted the proteomic analysis to proteins that were identified in all of the three TMT experiments. A total of N=4270 proteins passed the quality control filters. Log2-transformed raw TMT reporter ion intensities were first cleaned for batch effects using the “removeBatchEffects” function from limma v.3.54.2^64^ to minimize the influence of TMT batch on the intensity values. Next, batch corrected data was variance stabilization normalized using the vsn package v.3.66.0^65^. PCA was performed on the normalized TMT intensity data.

### Quality control of bulk-level datasets

To identify potential outliers, we performed sample clustering analysis in the miRNA-seq, RNA-seq, and proteomics datasets individually using principal component analysis (PCA). Here, we found two samples (CUD_3 and CUD_9) that consistently separated from the remaining samples across bulk-level datasets (Figure S1B). In a cell type deconvolution analysis on bulk RNA expression signatures using CIBERSORT (Figure S1C+D), we found that for these samples only a minimal proportion of medium spiny neurons (MSNs), the major neuronal cell type in the VS, was estimated. To minimize the influence of differential cell type proportion on the results of bulk-level analyses by excluding the two samples predicted to lack a major cell type of the VS, we restricted differential expression and downstream analyses including WGCNA and MOFA to the remaining N=38 samples.

### Differential microRNA expression analysis

Differential expression (DE) testing for the miRNA expression dataset was performed in DESeq2 v.1.38.3^66^ with covariate adjustment for donor age, PMI, brain pH, and RIN using the following DESeq2 model: *Expr(miRNA) ∼ CUD + age + PMI + pH + RIN*. Threshold criteria for differential miRNA expression in the CUD vs. Ctrl comparison were absolute log2FC larger than 0.07 (5% change in miRNA expression), an association with p<0.05 for nominal significance, and FDR-adjusted q<0.05 for significance after multiple testing correction. Principal component analysis (PCA) on miRNA-sequencing data was performed using variance stabilization transformed counts.

### microRNA target gene prediction

Target genes of nominally significant (p<0.05) miRNAs were predicted using miRNet2.0^29^ (https://www.mirnet.ca/upload/MirUploadView.xhtml) using miRTarBase v9.0, TarBase v9.0 and miRecords as the reference. We also included lncRNAs as potential miRNA target genes. Resulting “mir2gene” and “mir2lnc” files from miRNet were downloaded and filtered for target genes that were among nominally significant differentially expressed genes from RNA-seq analysis. This resulted in prioritized a list of miRNA targets that also show evidence for differential expression in CUD.

### Differential RNA expression analysis

Similar to the miRNA-sequencing dataset, DE testing on the raw count matrix in DESeq2 was performed using the same set of covariates in the statistical model: *Expr(RNA) ∼ CUD + age + PMI + pH + RIN*. Again, criteria for differential expression were absolute log2FC > 0.07, p<0.05 for nominal significance, and FDR-adjusted q<0.05 for transcriptome-wide significance. A PCA on RNA-seq expression data was performed using the variance stabilization transformed counts from DESeq2. To evaluate functional enrichment of DE results, a Gene Onotology (GO) enrichment analysis was performed based on transcripts with nominally significant association with CUD (p<0.05). The GO enrichment analysis was performed individually for up- and downregulated DE genes using the compareCluster function of the R package clusterProfiler v.4.6.2^67^. Terms with significant enrichment (FDR-adjusted q<0.05) were extracted from the GO enrichment results. For visualization, functional modules were generated using the emapplot function of enrichplot v.1.18.3.

### Differential expression analysis of snRNA-seq data

Annotation of clusters in the VS snRNA dataset was based on the expression of marker genes that have been previously used in other snRNA-seq studies of the brain^68,69^. Cell type specific DE analysis was performed in a CUD vs. Ctrl comparison using “FindMarkers” with min.pct=0.25 parameter to only test differential expression of genes that show robust expression levels in the cluster. DE testing was restricted to the N=7 major cell type clusters consisting of at least 100 cells from each condition (CUD/Ctrl). DE thresholds for the snRNA analysis were |log2FC| > 0.25 and p_adj_<0.05 (Bonferroni correction as implemented in Seurat). For visualization of DE results, a DE gene heatmap was generated using ComplexHeatmap v.2.14.0^70^ and the most significantly CUD-associated DEGs from each cluster were highlighted by gene name. Functional enrichment of cell type specific DEGs within biological pathways was evaluated using GO enrichment analysis. Significant results (FDR-adjusted q<0.05) were visualized in an enrichment map. DE gene overlap across evaluated clusters was analyzed in an upset plot created using UpSetR v.1.4.0^71^.

### Differential protein expression analysis

Differential expression testing for proteins in CUD cases and control individuals was performed based on the normalized, log2-transformed reporter intensities. The following statistical model was used in the model matrix that was given as an argument to the ‘lmFit’ function of limma: *Expr(protein) ∼ CUD + age + PMI + pH + batch (TMT)*. FDR-correction was used to adjust p-values for multiple testing. Protein DE cut-offs were the same as used in the miRNA-seq and RNA-seq analysis: |log2FC| > 0.07, p<0.05 for nominal significance, FDR-adjusted q<0.05 for proteome-wide significance. DE proteins associated with CUD at nominal significance were extracted for a GO enrichment analysis that was performed using clusterProfiler for upregulated and downregulated DE proteins individually.

### Cell type deconvolution analysis

Differences in the distribution of brain cell types across samples might lead to differential RNA expression independent of other phenotypes. We thus performed a cell type deconvolution analysis using CIBERSORT^72^ to evaluate the relationship between cell type percentage and differential expression results in the RNA-seq data. As we have generated single-nuclei RNA-seq data from a subset of the VS postmortem brain samples in this study, we created a customized gene expression reference matrix for the ventral striatum based on our snRNA data from N=16 individuals. From the snRNA dataset, cell type specific expression data was extracted for glial cells such as astrocytes, oligodendrocytes, oligodendrocyte precursor cells, and microglia. To obtain robust estimates, the different neuronal clusters were summarized to a medium-spiny neuron (MSN) cluster containing D1- and D2-MSNs (Inh_MSN) and an inhibitory GABA cluster (Inh_GABA) containing all other inhibitory, non-MSN neurons. Next, from the transcriptome-wide gene x cell normalized count matrix, we selected potential marker genes characterized by at least 10-times stronger expression in one cell type compared to all other cell types. The filtered gene expression matrix (3,081 genes in 20,492 cells) was used as the input dataset for the CIBERSORT.jar distribution resulting in a customized reference matrix for the VS. Using the CIBERSORT.R script (v.1.04, https://cibersortx.stanford.edu), CIBERSORT cell type estimates were generated for the bulk RNA-seq data in N=40 samples using normalized counts from DESeq2 (Figure S1C). To evaluate the accuracy of cell type estimation based on our customized reference matrix for the VS, we compared the measured cell type proportions as determined in snRNA-seq with the CIBERSORT estimates for the N=16 samples for which both bulk and snRNA-seq data are available. Pearson correlation coefficients ranged from *r*=0.44 to *r*=0.99 (median *r*=0.86) confirming successful generation of a customized reference matrix for the estimation of cell types in the VS (Figure S1D). Using the snRNA-seq derived reference matrix, differences in cell type proportion between CUD cases and control individuals were tested for the bulk RNA-seq dataset using the Bayesian estimation procedure from BEST^73^ (R package BayesianFirstAid v.0.1). Overlap between cell type markers derived from the snRNA-seq dataset with differential expression results from bulk RNA-seq was analyzed using GeneOverlap v.1.34.0^74^.

### Transcriptome-proteome correlation analysis

As few is known about the correlation between RNA and protein levels in postmortem human brain, the overall relationship between transcriptome and proteome was investigated using a correlation analysis approach. Following the strategy from Yang and Gorski ^75^, we performed preprocessing of transcriptomic and proteomic datasets. To make gene expression levels compatible with protein levels, the RNA-seq data was normalized to gene length and sequencing depth, and transcript per million (TPM) values were generated. For this, we performed pseudo-alignment of RNA-seq raw data to the GRCh38 primary assembly reference transcriptome using Salmon v.1.10.0^76^. From the Salmon quantification files, TPM estimates were extracted using tximport v.1.26.1^77^ followed by log2-transformation of TPM values. For the proteomic dataset we used the batch corrected, log2-transformed and vsn-normalized TMT intensity matrix. We performed filtering of RNA-seq and proteomics data for individuals that have both gene expression and protein data (N=39) and from the RNA-seq dataset we kept only protein coding genes that were detected in the proteomics dataset (N=3,935). At the sample level, within-individual correlation of transcriptome and proteome was determined using Pearson correlation. Next, we averaged RNA and protein expression levels across samples and calculated the correlation between mean RNA mean protein expression values for the N=3,935 genes. Further, Pearson correlation was assessed in a gene-centered approach resulting in a distribution of N=3,935 correlation coefficients. Based on the ranking of genes by correlation coefficients, we performed pre-ranked gene set enrichment analysis (GSEA) as implemented in clusterProfiler^67^ to evaluate functional enrichment of strongly positively and negatively correlated genes within biological pathways.

### WGCNA

To evaluate co-expression patterns in the miRNA-seq (N=1,542 miRNAs), RNA-seq (N= 22,685 RNAs), and proteomic datasets (N=4,270 proteins), a weighted correlation network analysis (WGCNA, package v.1.72.1)^78^ was performed. Using high-dimensional input datasets, WGCNA applies a pairwise correlation and hierarchical clustering approach to identify co-expression modules that are then related to trait variables such as the CUD phenotype. Normalized and variance stabilization transformed counts/TMT reporter intensities were used as the input datasets. Soft-power thresholds to achieve the criterion of scale-free topology (R^2^>0.9) were determined for each dataset individually by running the pickSoftThreshold function resulting in powers of 6, 7, and 9 for the miRNA-seq, RNA-seq and proteomics dataset, respectively. Next, automated network construction was performed with the parameters minModuleSize=10 for miRNA data, minModuleSize=20 for RNA and protein datasets, mergeCutHeight=0.15, and maxBlockSize=36,000. Pearson correlation coefficients of the module eigengene, corresponding to the first principal component of the module’s expression matrix, with trait data including CUD and other phenotypes such as pH, RIN, PMI, and CIBERSORT estimated cell type proportions, were determined in each dataset. Co-expression modules characterized by significant correlation with CUD (p<0.05) were characterized using GO enrichment analysis as implemented in the enrichGO function of clusterProfiler. Further, to evaluate the relationship between modules across datasets, module eigengene correlation was determined between CUD-associated modules in RNA and protein data using Pearson correlation. Finally, we performed an overlap analysis using a Fisher Test as implemented in GeneOverlap to evaluate potential enrichment of differentially expressed genes and proteins in CUD-associated co-expression modules.

### Multi-Omics Factor Analysis

An integrative multi-omic analysis of miRNA-seq, RNA-seq, and proteomic datasets was performed using Multi Omics Factor Analysis (MOFA)^79^. MOFA as implemented in the R package MOFA2 v.1.8.0 provides a statistical framework for high-dimensional (omics) data integration leveraging factor analysis for the unsupervised identification of lower-dimensional factor representations of the input datasets. Relationship of factors to trait variables such as CUD is assessed in downstream analyses allowing the identification of features (i.e. miRNAs, transcripts, or proteins) that show CUD-associated variability. During data preprocessing, normalized and variance stabilization transformed counts/TMT intensities were z-scaled using the mscale function from jyluMisc v.0.1.5 and the resulting matrices were used as the input datasets in MOFA. Due to the different dimensionalities of the omics datasets and its potential negative influence on MOFA model performance (N=1,542 miRNAs, N=22,685 RNAs, and N=4,270 proteins), the top 4,270 highly variable genes from RNA-seq data were filtered and thereby matched to the size of the proteomic dataset. Expression data for all N=1,542 miRNAs was included. In MOFA, default data and model options were used, whereas training options were modified using “convergence mode” – slow and a “drop_factor_threshold” of 0.01 to drop factors from the model than explain less than 1% variance in each view. The learned factors were inspected and factors significantly associated with CUD were investigated using GSEA based on the ranking of RNA and protein features by factor weights. GSEA was performed using the gseGO function of clusterProfiler.

### hdWGCNA

Cell type specific co-expression signatures were investigated using hdWGCNA^37^ (hdWGCNA, R package v.0.2.26, documentation from https://smorabit.github.io/hdWGCNA/index.html). hdWGCNA was performed based on the code implementation from its source publication (https://github.com/smorabit/hdWGCNA_paper). To ensure sufficient cluster sizes for robust module detection, D1-MSN and D2-MSN clusters were combined to an “Inh_MSN” cluster and inhibitory neuron clusters GABAergic-1, GABAergic-2, and GABAergic-3 were condensed to an “Inh_GABA” cluster.

Iteratively, hdWGCNA was performed in each cell type resulting in co-expression module labels according to cell types. Metacells were generated with nearest-neighbour k=25 followed by network construction using “TestSoftPowers” and “ConstructNetwork” functions with default parameters. Resulting co-expression modules for each cell type cluster were inspected for cluster-specific module eigengene expression using “ModuleFeaturePlot”. A co-expression module was considered cell-type specific if the module eigengene showed strongest expression in the cell type with the same name as the co-expression module and the module eigengene expression pattern was robust across the respective cell type cluster i.e. the association is not driven by a few cells only. Next, differential module eigengene testing was performed to identify co-expression modules that show significant differences (upregulation/downregulation) between CUD cases and Ctrl individuals. For further downstream analyses, we prioritized modules that i) showed strongest expression in the cell type cluster that was used for constructing the co-expression networks thereby addressing cell type specificity of modules and ii) displayed statistically significant module eigengene differences associated with CUD (p<0.05) indicating differential abundance of co-expression patterns in CUD (Table S12). In the resulting module subset, pathway enrichment analyses were performed using the “RunEnrichr” function in hdWGCNA using GO and KEGG databases as reference. Results were ranked by the combined.score metric from Enrichr defined as log(p) from Fisher-Test multiplied by the z-score as the deviation from the expected rank. Using GeneOverlap, we tested the enrichment of cluster-specific CUD-associated DE genes within co-expression modules.

### Consensus hdWGCNA

As an integrative approach, we performed a cross-species consensus network analysis in hdWGCNA based on publicly available snRNA-seq data from a repeated cocaine-exposure model in rats ^25^. Sequencing data for the nucleus accumbens from male rats was downloaded from GEO (accession number: GSE222418) and processed using the same analysis pipeline as described for the analysis of the human dataset. Clusters in the rat dataset were annotated based on cell type markers from the original publication and neuronal subclusters were condensed into Inh_MSN and Inh_GABA clusters as previously described for the human dataset. Network construction was performed on expression data for homologous genes across species and from the identified consensus modules, we again selected modules that showed i) strongest module gene expression in the cell type of interest and ii) significant differential module eigengene association with CUD (Table S15). The overlap of module genes between CUD-associated DME modules from the analysis in human CUD and the consensus co-expression modules across species was evaluated using GeneOverlap. Finally, pathway enrichment analysis was performed in the CUD-associated consensus modules that most strongly overlapped between the human and consensus analyses.

### CellChat

We applied CellChat^80^ (R package v.2.1.2) in our snRNA-seq dataset to analyze CUD-associated expression changes of ligand-receptor pairs in MSNs and astrocytes. Following the documentation for combined analysis of multiple datasets (https://github.com/jinworks/CellChat/blob/main/tutorial/Comparison_analysis_of_multiple_datasets.html), construction of CellChat objects was performed individually in CUD and Ctrl nuclei using the human reference dataset for ligand-receptor interaction (CellChatDB.human). Next, the individual objects were merged using “mergeCellChat”. A differential expression analysis for ligand receptor (LR) pairs was performed using “identifyOverExpressedGenes” and “netMappingDEG” to identify ligands and receptors with statistically significant deregulation in CUD using tresh.pc=0.1, thresh.fc=0, and thresh.p=0.05 cut-offs. Significant up- and downregulated LR pairs were filtered for interactions with a minimum of 5% change in ligand expression (|log2FC>0.07|) and an expression change of its receptor into the same direction. The “netVisual_chord_gene” function was used to visualize differential LR interactions individually for up- and downregulated LR pairs in CUD. A donut plot was generated to summarize the distribution of signaling type annotations among up- and downregulated LR pairs.

## Acknowledgements

Funding supporting this study was provided by the German Federal Ministry of Education and Research (BMBF) within the e:Med research program SysMedSUDs: “A systems-medicine approach toward distinct and shared resilience and pathological mechanisms of substance use disorders” (01ZX01909 to R.S., P.K., M.R., A.C.H., and S.H.W.). In addition, by the Deutsche Forschungsgemeinschaft (DFG) through the collaborative research centre TRR265: “Losing and Regaining Control over Drug Intake”^52,81^ (Project ID 402170461 to S.H.W., R.S., A.C.H. and M.R.), the Hetzler Foundation for Addiction Research (to A.C.H.), and the ERA-NET program: Psi-Alc (01EZ1908). The project has been carried out using the Mannheim (CIMH) infrastructure of the German Center for Mental Health (DZPG). We thank Dr. Jennifer Schwarz and Dr. Frank Stein from the Proteomics Core Facility at the European Molecular Biology Laboratory (EMBL, Heidelberg, Germany) for their support in proteomic analyses. We further thank Elisabeth Röbel and Claudia Schäfer-Arnold for their technical assistance.

## Author contributions

Conceptualization, E.Z., L.Z., M.R., R.S., and S.H.W.; Methodology, E.Z., A.A., M.M.N, A.C.R., C.C.W., J.F., and L.Z.; Resources, G.T., N.M., P.K. and A.C.H, Data curation, E.Z., A.A., A.C.R., D.A., H.B., and L.Z. ; Data Analysis, E.Z., D.A., H.B., and L.Z., Investigation, E.Z., A.A., A.C.R., D.A., H.B., A.C.H., M.R., R.S., S.H.W, and L.Z.; Writing – Original Draft, E.Z., Writing – Reviewing & Editing, E.Z., A.A., A.C.R., D.A., H.B., J.F., N.M., G.T., M.M.N, A.C.H., C.C.W., M.R., P.K., R.S., S.H.W., and L.Z.; Supervision, J.F., M.R., C.C.W., P.K., R.S., S.H.W, and L.Z., Project Administration, M.M.N, P.K., R.S., A.C.H., M.R. and S.H.W., Funding Acquisition, P.K., R.S., A.C.H., M.R., and S.H.W.

## Declaration of interests

The authors declare that there are no competing interests.

## Notes

### Competing Interest Statement

The authors have declared no competing interest.

