## Supplementary Figures for "A multi-omics and cell type-specific characterization of the ventral striatum in human cocaine use disorder"

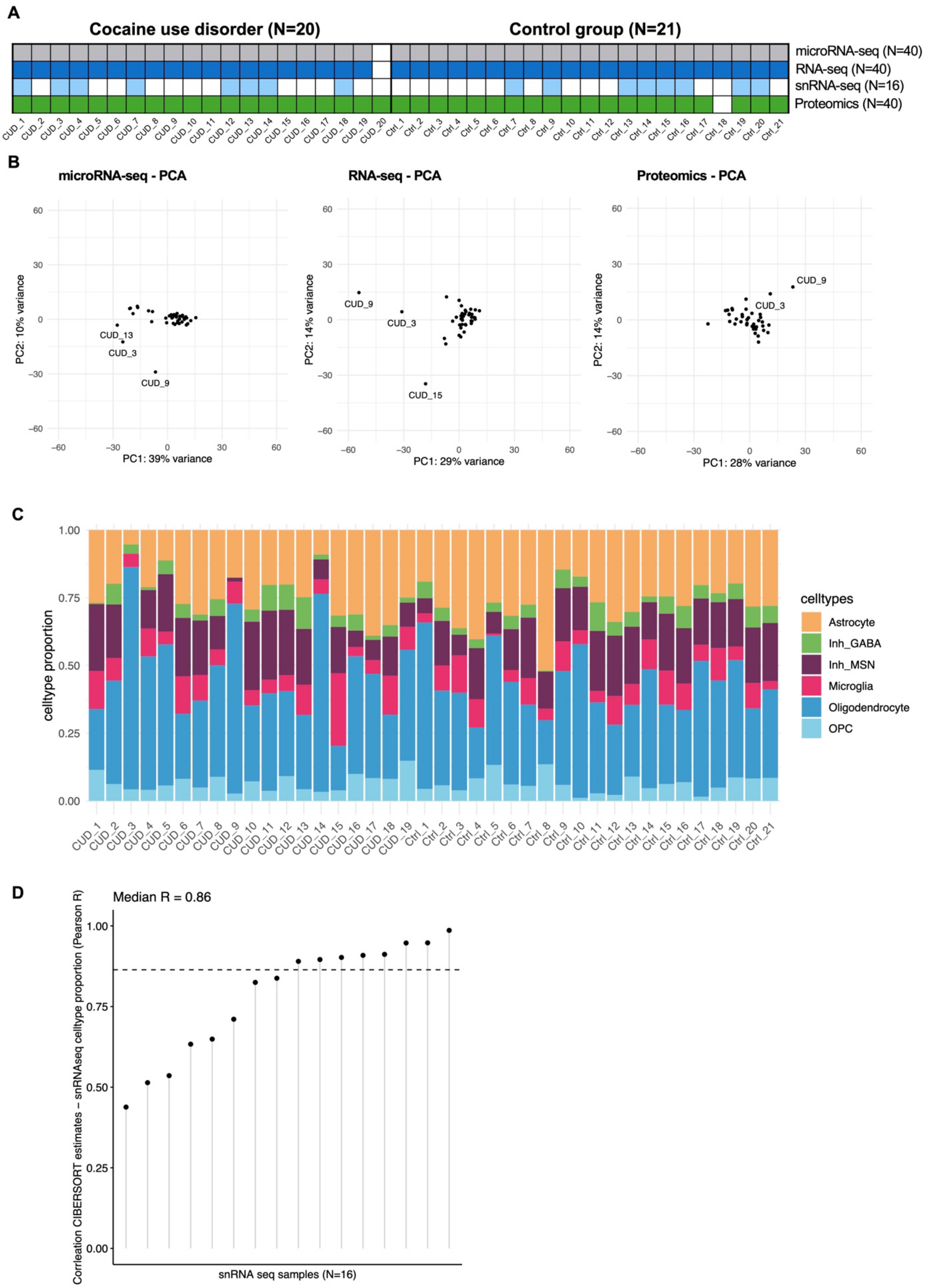
A
Cocaine use disorder (N=20)
Control group (N=21)
microRNA-seq (N=40)
RNA-seq (N=40)
snRNA-seq (N=16)
Proteomics (N=40)
CUD_1 CUD_2 CUD_3 CUD_4 CUD_5 CUD_6 CUD_7 CUD_8 CUD_9 CUD_10 CUD_11 CUD_12 CUD_13 CUD_14 CUD_15 CUD_16 CUD_17 CUD_18 CUD_19 CUD_20
Ctrl_1 Ctrl_2 Ctrl_3 Ctrl_4 Ctrl_5 Ctrl_6 Ctrl_7 Ctrl_8 Ctrl_9 Ctrl_10 Ctrl_11 Ctrl_12 Ctrl_13 Ctrl_14 Ctrl_15 Ctrl_16 Ctrl_17 Ctrl_18 Ctrl_19 Ctrl_20 Ctrl_21
B
microRNA-seq - PCA
PC2: 10% variance
PC1: 39% variance
CUD_13
CUD_3
CUD_9
RNA-seq - PCA
PC2: 14% variance
PC1: 29% variance
CUD_9
CUD_3
CUD_15
Proteomics - PCA
PC2: 14% variance
PC1: 28% variance
CUD_9
CUD_3
C
celltype proportion
celltypes
Astrocyte
Inh_GABA
Inh_MSN
Microglia
Oligodendrocyte
OPC
D
Median R = 0.86
Corrleation CIBERSORT estimates ~ snRNAseq celltype proportion (Pearson R)
snRNA seq samples (N=16)

**Supplementary Figure 1 – Bulk dataset overview and cell type deconvolution analysis**

**A** comparative overview on dataset availability in the N=41 postmortem brain tissue samples of the ventral striatum (VS). **B** inspection of bulk datasets by principal component analysis in miRNA-seq, RNA-seq and proteomic datasets. **C** results of a cell type deconvolution analysis (CIBERSORT) for bulk RNA-seq data indicating estimated proportions of cell types in bulk RNA-seq samples. The reference dataset for the VS was generated using cell type specific expression information from the N=16 snRNA-seq samples. Cell type deconvolution analysis was restricted to major cell type of the VS where DRD1- and DRD2-expressing medium spiny neurons (MSMs) were assigned to an Inh_MSN category and other non-MSN GABAergic interneurons were condensed to an Inh_GABA category. **D** correlation analysis between true cell type proportions identified by snRNA-seq and inferred proportions from CIBERSORT for the N=16 VS samples for which both bulk RNA-seq and snRNA-seq data was available.

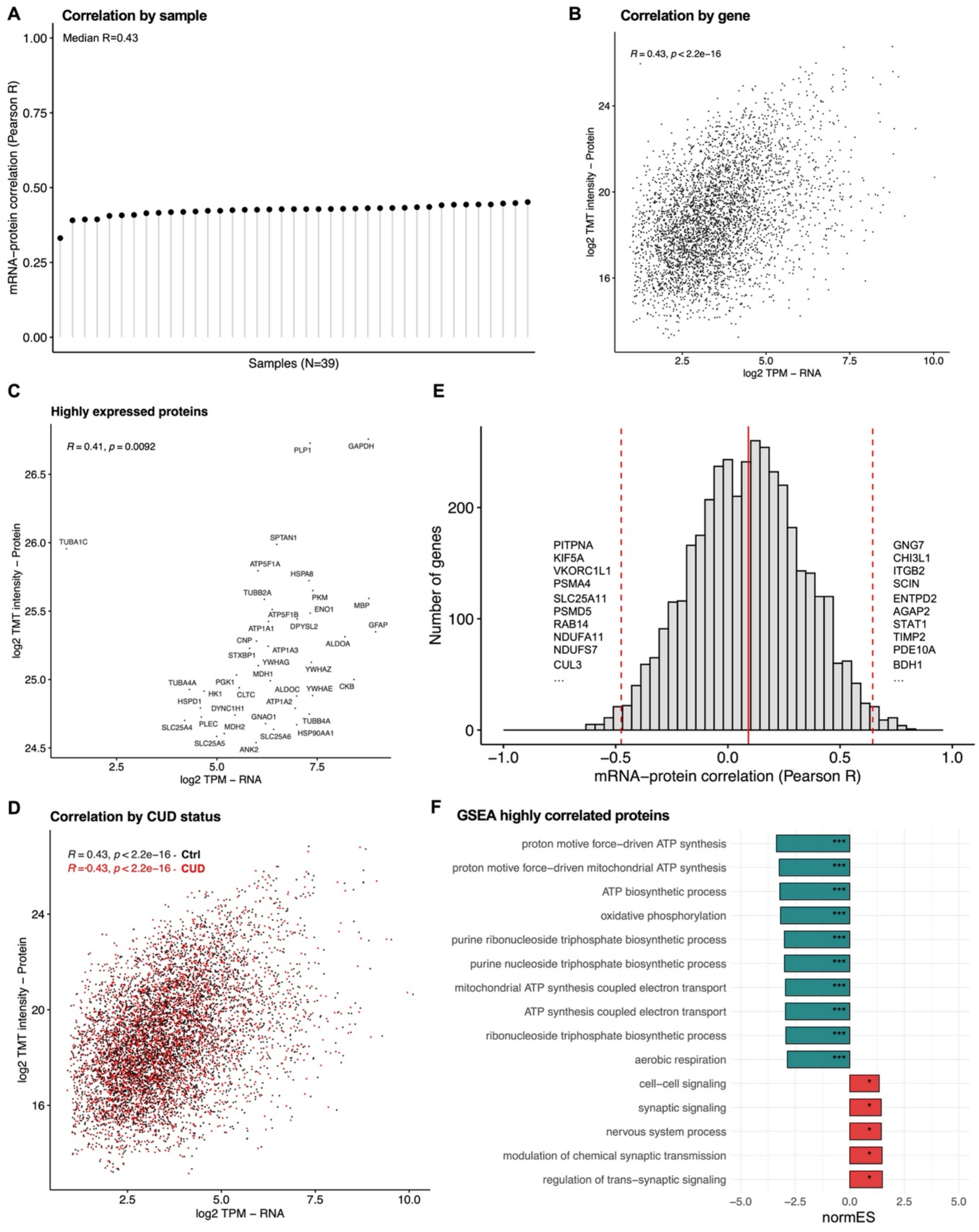
A
Correlation by sample
Median R=0.43
mRNA−protein correlation (Pearson R)
1.00
0.75
0.50
0.25
0.00
Samples (N=39)
B
Correlation by gene
R = 0.43, p < 2.2e−16
log2 TMT intensity − Protein
24
20
16
2.5
5.0
7.5
10.0
log2 TPM − RNA
C
Highly expressed proteins
R = 0.41, p = 0.0092
log2 TMT intensity − Protein
26.5
26.0
25.5
25.0
24.5
2.5
5.0
7.5
log2 TPM − RNA
PLP1
GAPDH
TUBA1C
SPTAN1
ATP5F1A
HSPA8
TUBB2A
PKM
ENO1
MBP
ATP5F1B
ATP1A1
DPYSL2
GFAP
ALDOA
CNP
ATP1A3
STXBP1
YWHAG
YWHAZ
MDH1
TUBA4A
PGK1
CKB
HK1
CLTC
ALDOC
YWHAE
HSPD1
DYNC1H1
ATP1A2
GNAO1
TUBB4A
SLC25A4
PLEC
MDH2
HSP90AA1
SLC25A6
SLC25A5
ANK2
D
Correlation by CUD status
R = 0.43, p < 2.2e−16 - Ctrl
R = 0.43, p < 2.2e−16 - CUD
log2 TMT intensity − Protein
24
20
16
2.5
5.0
7.5
10.0
log2 TPM − RNA
E
Number of genes
200
100
0
−1.0
−0.5
0.0
0.5
1.0
mRNA−protein correlation (Pearson R)
PITPNA
KIF5A
VKORC1L1
PSMA4
SLC25A11
PSMD5
RAB14
NDUFA11
NDUFS7
CUL3
...
GNG7
CHI3L1
ITGB2
SCIN
ENTPD2
AGAP2
STAT1
TIMP2
PDE10A
BDH1
...
F
GSEA highly correlated proteins
proton motive force−driven ATP synthesis
proton motive force−driven mitochondrial ATP synthesis
ATP biosynthetic process
oxidative phosphorylation
purine ribonucleoside triphosphate biosynthetic process
purine nucleoside triphosphate biosynthetic process
mitochondrial ATP synthesis coupled electron transport
ATP synthesis coupled electron transport
ribonucleoside triphosphate biosynthetic process
aerobic respiration
cell−cell signaling
synaptic signaling
nervous system process
modulation of chemical synaptic transmission
regulation of trans−synaptic signaling
***
*
−5.0
−2.5
0.0
2.5
5.0
normES

**Supplementary Figure 2 - Correlation analysis between transcriptome and proteome in the ventral striatum**

**A** sample-level correlation analysis between RNA and protein expression levels of N=3,935 genes. The analysis was performed in N=39 ventral striatum samples for which both transcriptomic and proteomic data was available. **B** correlation analysis of mean RNA (log2-transcript per million, log2 TPM) and protein expression values (log2 TMT-intensities) for each of the N=3,935 genes across samples. **C** correlation analysis of mean expression levels for RNA and proteins after selecting for highly expressed proteins ($99^{th}$ percentile of log2 TMT intensities). **D** sensitivity analysis of the correlation approach separated by CUD status. **E** histogram of Pearson correlation coefficients from a gene-centered approach suggesting genes with strong positive (right) and negative (left) correlation of RNA and protein expression levels. RNA and protein expression levels were correlated individually for each gene without prior averaging of expression levels across samples. **F** pre-ranked gene-set enrichment analysis (GSEA) using gene-level correlation coefficients from E as the ranking metric. GSEA results separated by normalized enrichment scores (normES) suggest pathways with statistically significant enrichment for genes with strong positive (red) and negative correlation between RNA and protein expression levels. *$p<0.05$, **$p<0.01$, ***$p<0.001$.

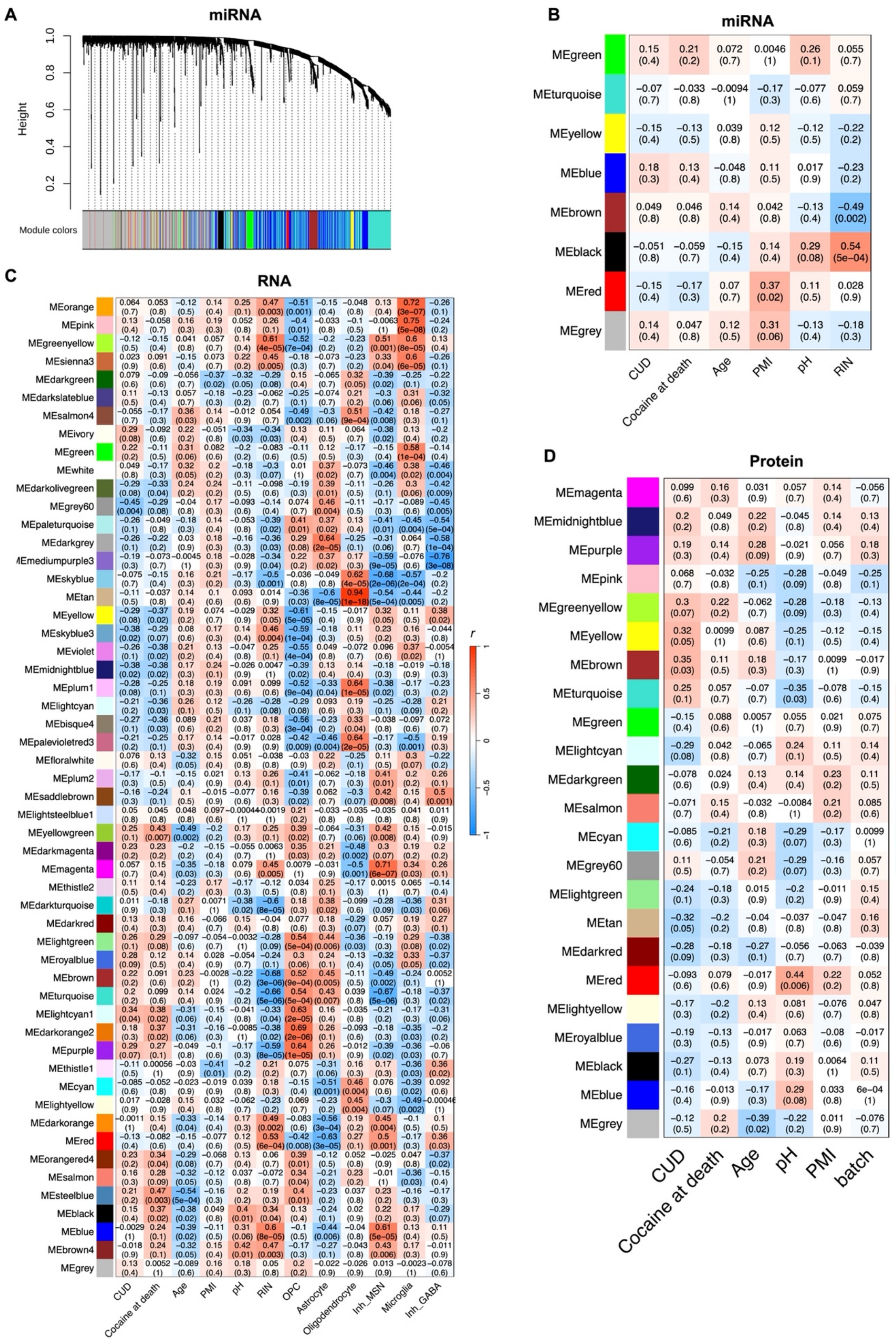
A
miRNA
Height
Module colors
B
miRNA
MEgreen
MEturquoise
MEyellow
MEblue
MEbrown
MEblack
MEred
MEgrey
CUD
Cocaine at death
Age
PMI
pH
RIN
r
C
RNA
MEorange
MEpink
MEgreenyellow
MEsienna3
MEdarkgreen
MEdarkslateblue
MEsalmon4
MEivory
MEgreen
MEwhite
MEdarkolivegreen
MEgrey60
MEpaleturquoise
MEdarkgrey
MEmediumpurple3
MEskyblue
MEtan
MEyellow
MEskyblue3
MEviolet
MEmidnightblue
MEplum1
MElightcyan
MEbisque4
MEpalevioletred3
MEfloralwhite
MEplum2
MEsaddlebrown
MElightsteelblue1
MEyellowgreen
MEdarkmagenta
MEmagenta
MEthistle2
MEdarkturquoise
MEdarkred
MElightgreen
MEroyalblue
MEbrown
MEturquoise
MElightcyan1
MEdarkorange2
MEpurple
MEthistle1
MEcyan
MElightyellow
MEdarkorange
MEred
MEorangered4
MEsalmon
MEsteelblue
MEblack
MEblue
MEbrown4
MEgrey
CUD
Cocaine at death
Age
PMI
pH
RIN
OPC
Astrocyte
Oligodendrocyte
Inh_MSN
Microglia
Inh_GABA
D
Protein
MEmagenta
MEmidnightblue
MEpurple
MEpink
MEgreenyellow
MEyellow
MEbrown
MEturquoise
MEgreen
MElightcyan
MEdarkgreen
MEsalmon
MEcyan
MEgrey60
MElightgreen
MEtan
MEdarkred
MEred
MElightyellow
MEroyalblue
MEblack
MEblue
MEgrey
CUD
Cocaine at death
Age
pH
PMI
batch

**Supplementary Figure 3 - Weighted correlation network analysis (WGCNA) in bulk-level miRNA-seq, RNA-seq, and proteomic datasets of the ventral striatum in cocaine use disorder**

**A** dendrogram of the miRNA-seq co-expression network analysis using expression information. **B** correlation analysis of miRNA co-expression module eigengenes (ME) with covariates including cocaine use disorder (CUD) and other available phenotypic variables. Panels show Pearson correlation coefficients (*r*) with correlation p-value in brackets. PMI: postmortem interval, pH: postmortem brain tissue pH value, RIN: RNA integrity number. **C** correlation of MEs from RNA co-expression modules with phenotypic variables and cell type estimates from deconvolution analysis in CIBERSORT. OPC: oligodendrocyte progenitor cell, Inh_MSN: cell type estimates for DRD1- and DRD2-expressing medium spiny neurons (MSNs), Inh_GABA: cell type estimates for GABAergic, non-MSN interneurons. **D** correlation of MEs from protein co-expression modules with phenotypic information. batch: proteomics processing batch.

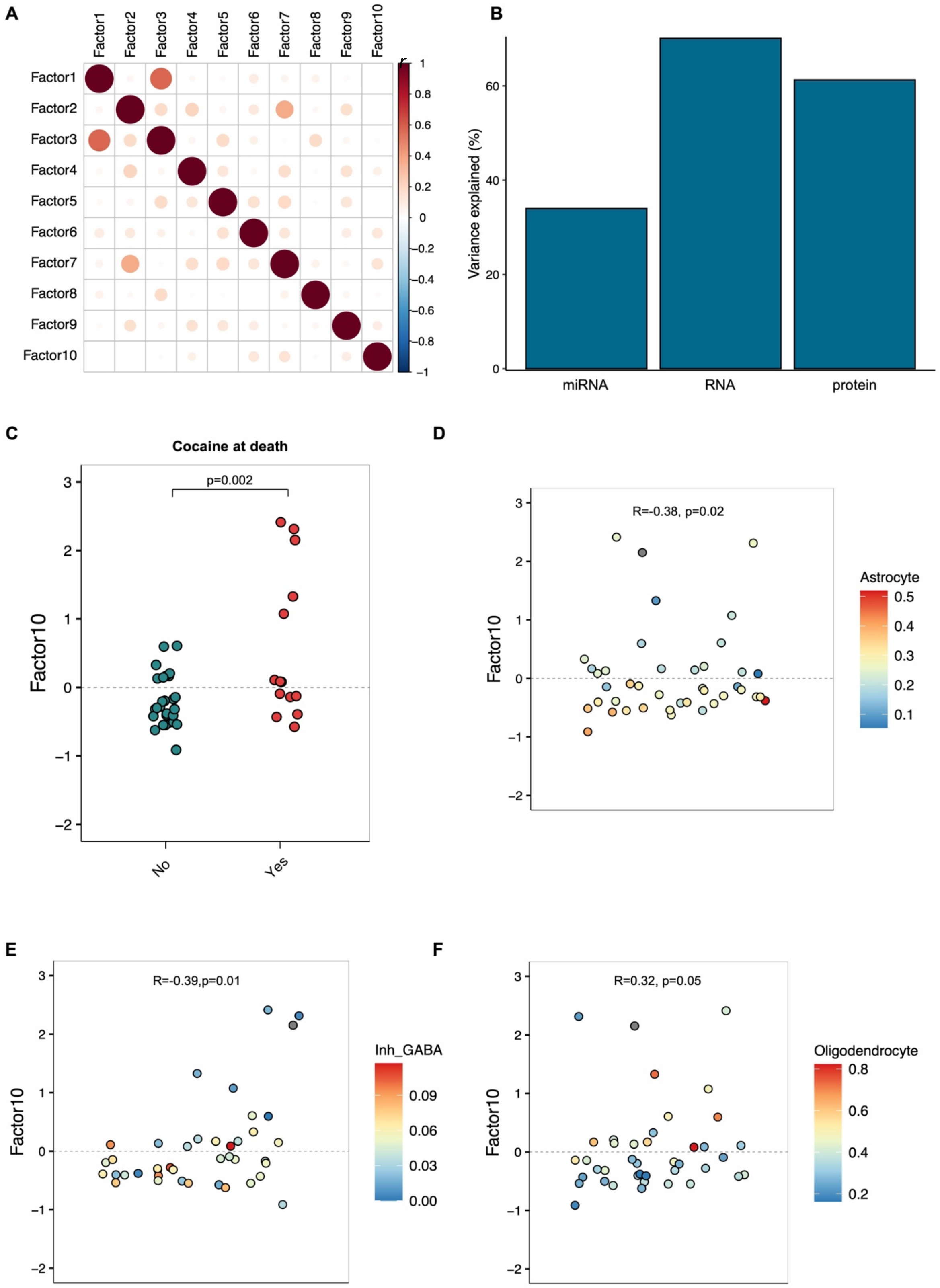
A
Factor1
Factor2
Factor3
Factor4
Factor5
Factor6
Factor7
Factor8
Factor9
Factor10
1
0.8
0.6
0.4
0.2
0
−0.2
−0.4
−0.6
−0.8
−1
B
Variance explained (%)
0
20
40
60
miRNA
RNA
protein
C
Cocaine at death
p=0.002
Factor10
No
Yes
D
R=-0.38, p=0.02
Astrocyte
0.5
0.4
0.3
0.2
0.1
E
R=-0.39,p=0.01
Inh_GABA
0.09
0.06
0.03
0.00
F
R=0.32, p=0.05
Oligodendrocyte
0.8
0.6
0.4
0.2

**Supplementary Figure 4 – Characterization of a cocaine use disorder associated factor identified by Multi-Omics Factor Analysis (MOFA)**

**A** intercorrelation analysis of multi-omics dataset latent factor representations using dot representation of Pearson correlation coefficients (*r*). **B** total variance explanation in miRNA-seq (miRNA), RNA-seq (RNA) and proteomics (protein) datasets by the learned factor model from MOFA. **C** association of MOFA factor 10 with cocaine at death status (yes/no). p-value from Wilcoxon Rank-sum test. Statistically significant correlation of factor 10 values with cell type estimates from CIBERSORT for **D** astrocyte, **E** oligodendrocyte, and **F** Inh_GABA cell type categories.

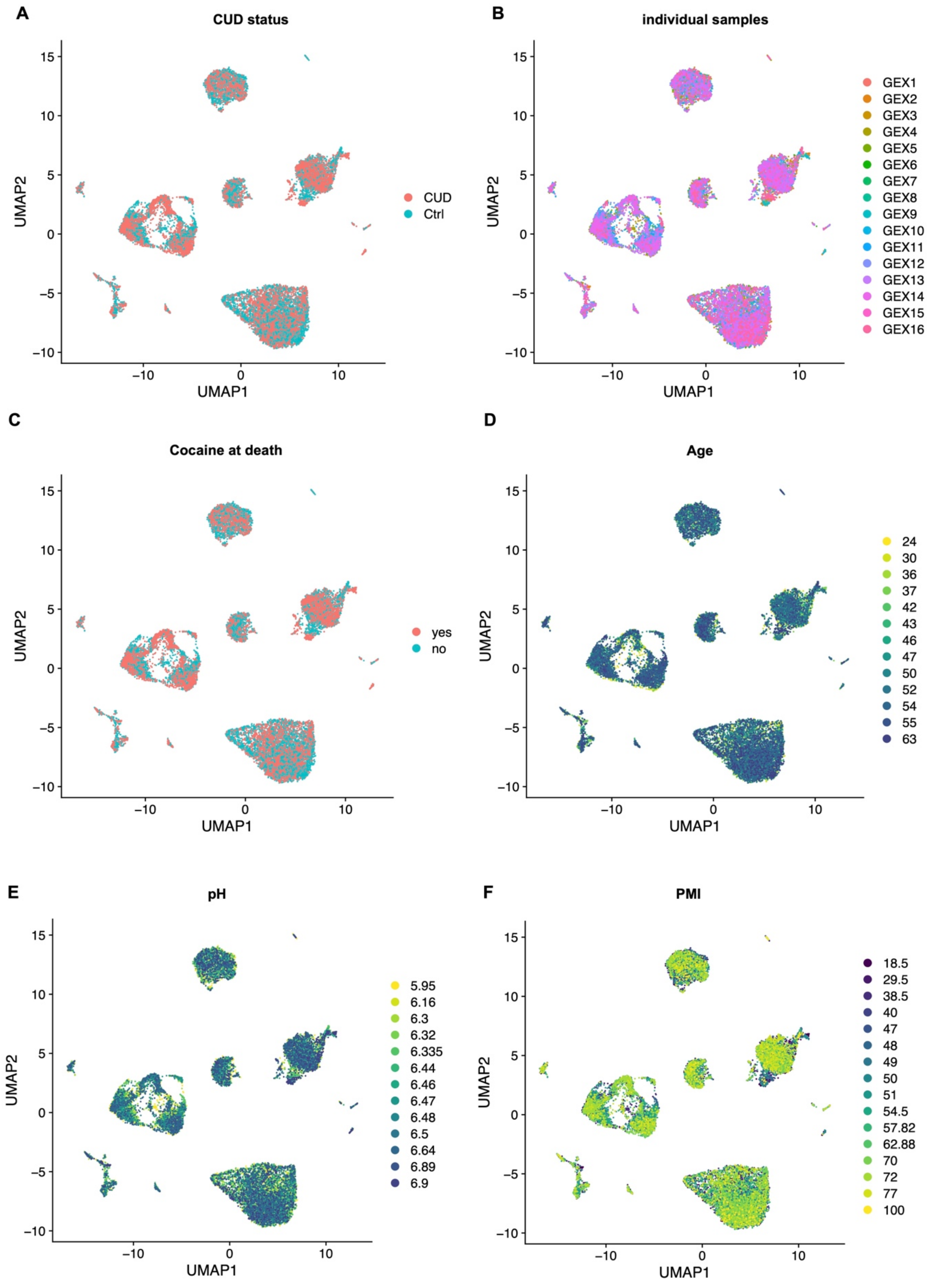
A
CUD status
UMAP2
UMAP1
CUD
Ctrl
B
individual samples
GEX1
GEX2
GEX3
GEX4
GEX5
GEX6
GEX7
GEX8
GEX9
GEX10
GEX11
GEX12
GEX13
GEX14
GEX15
GEX16
C
Cocaine at death
yes
no
D
Age
24
30
36
37
42
43
46
47
50
52
54
55
63
E
pH
5.95
6.16
6.3
6.32
6.335
6.44
6.46
6.47
6.48
6.5
6.64
6.89
6.9
F
PMI
18.5
29.5
38.5
40
47
48
49
50
51
54.5
57.82
62.88
70
72
77
100

**Supplementary Figure 5 – UMAP representation of cell type specific RNA expression profiles in N=20,759 single nuclei of the ventral striatum in cocaine use disorder**

For the 12 distinct cell type clusters identified by single-nuclei (sn)RNA-seq in N=16 VS samples, the association of clustering patterns with phenotypes was assessed. **A** CUD status (CUD/Ctrl), **B** N=16 snRNA-seq samples (GEX1-GEX16), **C** cocaine at death status (yes/no), **D** tissue donor age (years), **E** brain tissue pH value, **F** postmortem interval (hours).

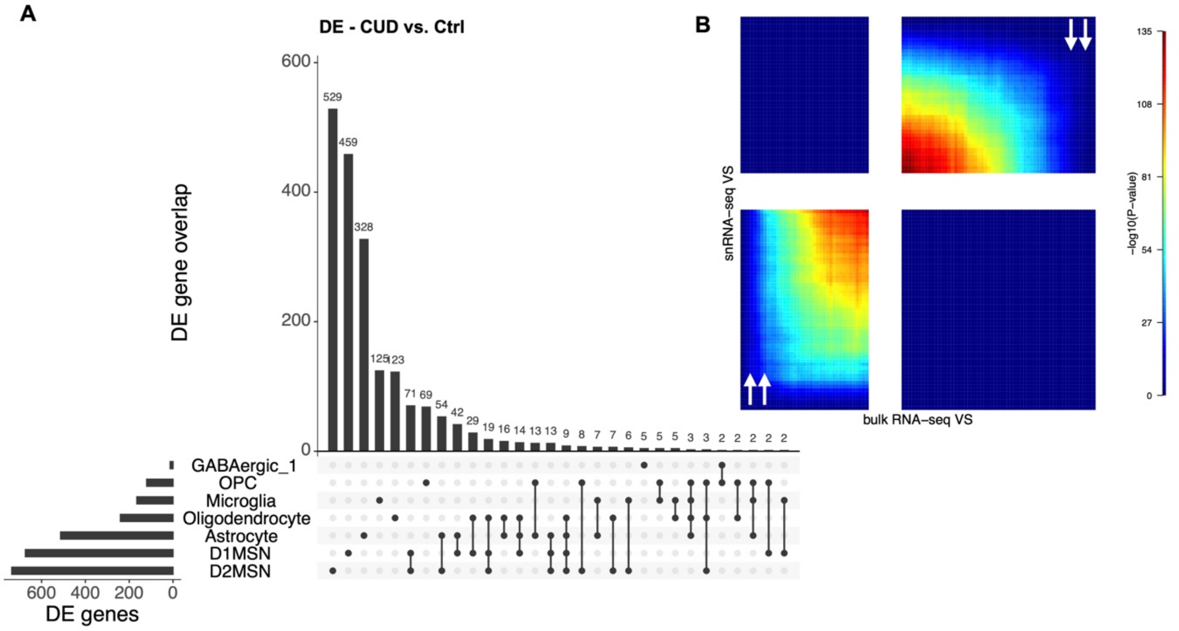


**Supplementary Figure 6 – Comparison of cell type specific differential expression profiles associated with cocaine use disorder**

**A** overlap between the N=1,996 differentially expressed genes (|log2FC| > 0.25, FDR q<0.05) from 7 major cell types of the VS was visualized in an upset plot. **B** rank-rank hypergeometric overlap analysis of CUD-associated differential expression patterns from i) bulk RNA-seq in N=38 samples and ii) snRNA-seq in N=16 samples of the ventral striatum (VS). Arrows in RRHO panels indicate convergent upregulation (bottom-left) or downregulation (top-right) of expression patterns across datasets.

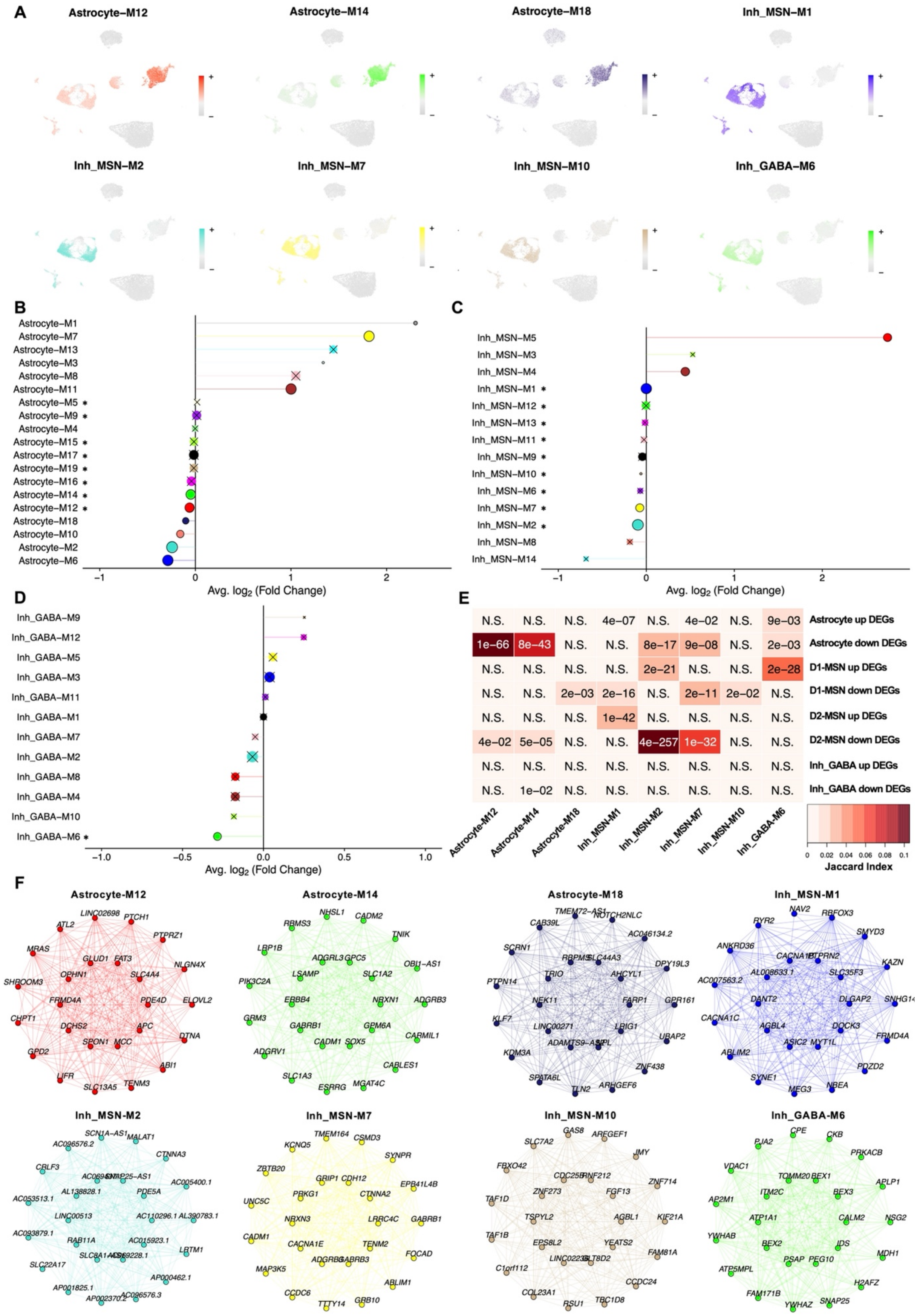
A
Astrocyte–M12
Astrocyte–M14
Astrocyte–M18
Inh_MSN–M1
Inh_MSN–M2
Inh_MSN–M7
Inh_MSN–M10
Inh_GABA–M6
B
Astrocyte–M1
Astrocyte–M7
Astrocyte–M13
Astrocyte–M3
Astrocyte–M8
Astrocyte–M11
Astrocyte–M5 *
Astrocyte–M9 *
Astrocyte–M4
Astrocyte–M15 *
Astrocyte–M17 *
Astrocyte–M19 *
Astrocyte–M16 *
Astrocyte–M14 *
Astrocyte–M12 *
Astrocyte–M18
Astrocyte–M10
Astrocyte–M2
Astrocyte–M6
Avg. log2 (Fold Change)
C
Inh_MSN–M5
Inh_MSN–M3
Inh_MSN–M4
Inh_MSN–M1 *
Inh_MSN–M12 *
Inh_MSN–M13 *
Inh_MSN–M11 *
Inh_MSN–M9 *
Inh_MSN–M10 *
Inh_MSN–M6 *
Inh_MSN–M7 *
Inh_MSN–M2 *
Inh_MSN–M8
Inh_MSN–M14
Avg. log2 (Fold Change)
D
Inh_GABA–M9
Inh_GABA–M12
Inh_GABA–M5
Inh_GABA–M3
Inh_GABA–M11
Inh_GABA–M1
Inh_GABA–M7
Inh_GABA–M2
Inh_GABA–M8
Inh_GABA–M4
Inh_GABA–M10
Inh_GABA–M6 *
Avg. log2 (Fold Change)
E
Astrocyte up DEGs: N.S. N.S. N.S. 4e–07 N.S. 4e–02 N.S. 9e–03
Astrocyte down DEGs: 1e–66 8e–43 N.S. N.S. 8e–17 9e–08 N.S. 2e–03
D1-MSN up DEGs: N.S. N.S. N.S. N.S. 2e–21 N.S. N.S. 2e–28
D1-MSN down DEGs: N.S. N.S. 2e–03 2e–16 N.S. 2e–11 2e–02 N.S.
D2-MSN up DEGs: N.S. N.S. N.S. 1e–42 N.S. N.S. N.S. N.S.
D2-MSN down DEGs: 4e–02 5e–05 N.S. N.S. 4e–257 1e–32 N.S. N.S.
Inh_GABA up DEGs: N.S. N.S. N.S. N.S. N.S. N.S. N.S. N.S.
Inh_GABA down DEGs: N.S. 1e–02 N.S. N.S. N.S. N.S. N.S. N.S.
Astrocyte–M12
Astrocyte–M14
Astrocyte–M18
Inh_MSN–M1
Inh_MSN–M2
Inh_MSN–M7
Inh_MSN–M10
Inh_GABA–M6
Jaccard Index
F
Astrocyte–M12
Astrocyte–M14
Astrocyte–M18
Inh_MSN–M1
Inh_MSN–M2
Inh_MSN–M7
Inh_MSN–M10
Inh_GABA–M6

**Supplementary Figure 7 – Characterization of cell type specific co-expression networks identifies medium spiny neuron- and astrocyte-specific modules in cocaine use disorder**
Co-expression networks in the snRNA-seq dataset were identified for major cell types using hdWGCNA. **A** module eigengene expression of cell type specific co-expression modules in CUD-associated astrocyte, Inh_GABA, and Inh_MSN clusters. Differential module eigengene expression analysis in CUD for **B** astrocytes, **C** medium spiny neurons (MSN, D1+D2 combined), and **D** non-MSN inhibitory GABAergic neurons (Inh_GABA). Co-expression modules were prioritized based on significant DME in the CUD vs. Ctrl comparison (colored dots with X) and cell type specificity (*) measured by module eigengene expression in the respective cell types. **E** overlap analysis between module genes from CUD-associated cell type specific co-expression modules and differentially expressed (DE) genes from cluster-specific DE analysis. Jaccard index and p-values from Fisher-Test are shown. N.S.: not significant. **F** network plots of top 25 co-expression module hub genes for all cell type specific co-expression modules with significant DME in CUD.

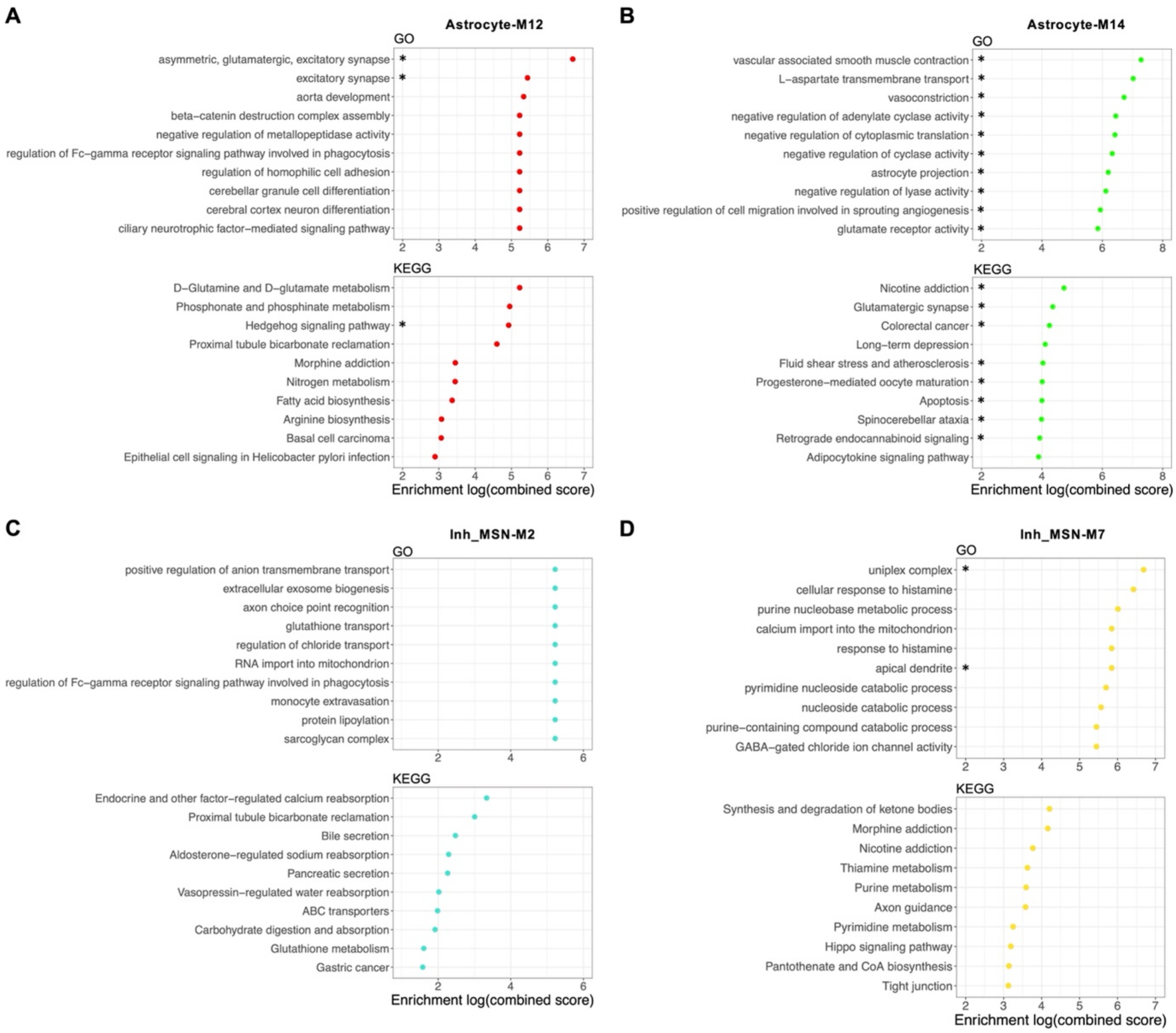


**Supplementary Figure 8 – Pathway enrichment analysis in astrocyte- and neuron-specific co-expression modules identifies altered glutamatergic and cAMP signaling in astrocytes and metabolic changes in medium spiny neurons**

Results of GO and KEGG pathway enrichment analyses in co-expression modules **A** Astrocyte-M12, **B** Astrocyte-M14, **C** Inh_MSN-M2, and **D** Inh_MSN-M7. These co-expression modules were characterized by strongest enrichment of module genes in cell type specific DEGs (see Figure S7E). The combined score metrics for results ranking was derived from the enrichment statistics in EnrichR, *FDR q<0.05.
